# A molecular atlas of the human postmenopausal fallopian tube and ovary from single-cell RNA and ATAC sequencing

**DOI:** 10.1101/2022.08.04.502826

**Authors:** Ernst Lengyel, Yan Li, Melanie Weigert, Lisha Zhu, Heather Eckart, Melissa Javellana, Sarah Ackroyd, Jason Xiao, Susan Olalekan, Dianne Glass, Shilpa Iyer, Agnes Julia Bilecz, Ricardo Lastra, Mengjie Chen, Anindita Basu

**Author notes:** These authors contributed equally: A. Basu, M. Chen, Y. Li, E. Lengyel.

## Abstract

As part of the Human Cell Atlas initiative, we generated transcriptomic (scRNA-seq; 86,708 cells) and regulatory (scATAC-seq; 59,118 cells) profiles of the normal postmenopausal ovary and fallopian tube (FT) at single-cell resolution. In the FT, 22 cell clusters integrated into 11 cell types, including ciliated and secretory epithelial cells, while the ovary had 17 distinct cell clusters defining 6 major cell types. The dominant cell type in both the postmenopausal ovary and FT was stromal cells, which expressed several genes associated with aging. The fimbrial end of the FT had a significant number and variety of immune cells and active communication with the ovary; the ovary contained mostly stromal cells but few immune cells. The epithelial cells of the normal FT expressed multiple ovarian cancer risk-associated genes (*CCDC170, RND3, TACC2, STK33*, and *ADGB*). By integrating paired single-cell transcriptomics and chromatin accessibility data we found that the regulatory landscape of the fimbriae was markedly different from the isthmus and ampulla. Intriguingly, several cell types in the FT had comparable gene expression but different transcriptional regulations. Our single-cell transcriptional and regulatory maps allowed us to disentangle the complex cellular makeup of the postmenopausal FT and ovary and will contribute to a better understanding of gynecologic diseases in menopause.

## Introduction

Several diseases of the fallopian tube and ovary, which cumulatively affect many women, can only be treated symptomatically because their pathophysiology is poorly understood, and their cells of origin are largely unknown. The first step towards a better understanding of the etiology of these tubal and ovarian diseases, which often surface after menopause, is to create a comprehensive map of the normal anatomy and cellular compositions of these organs. The three main areas of the fallopian tube, which transports the ovum to the uterus, explored in this study are 1) the isthmus, a short section with a thick muscular wall that is closest to the uterus and merges with 2) the ampulla, a longer, thinner-walled central portion, which connects to 3) the fimbriated end, shaped like the bell of a trumpet and fringed with fimbria, which opens and is in close contact with the ovary (**Fig. 1A**). The entire tube is composed of three layers. The innermost is an epithelial/mucosal layer lining the central muscular layer, consisting mostly of smooth muscle cells, and lines the third layer, an outer surface covered by serosa. There are no major histological differences between the pre- and postmenopausal fallopian tube. In contrast, the postmenopausal ovary differs dramatically from the premenopausal ovary in that it is mostly fibrotic with atretic follicles, covered by a single layer of cuboidal epithelial cells. During the reproductive years, the biological functions of the ovary include hormone production, oocyte maturation, and immune defense^1,2^, but its postmenopausal functions and cellular compositions are less well defined.

**Figure 1:**
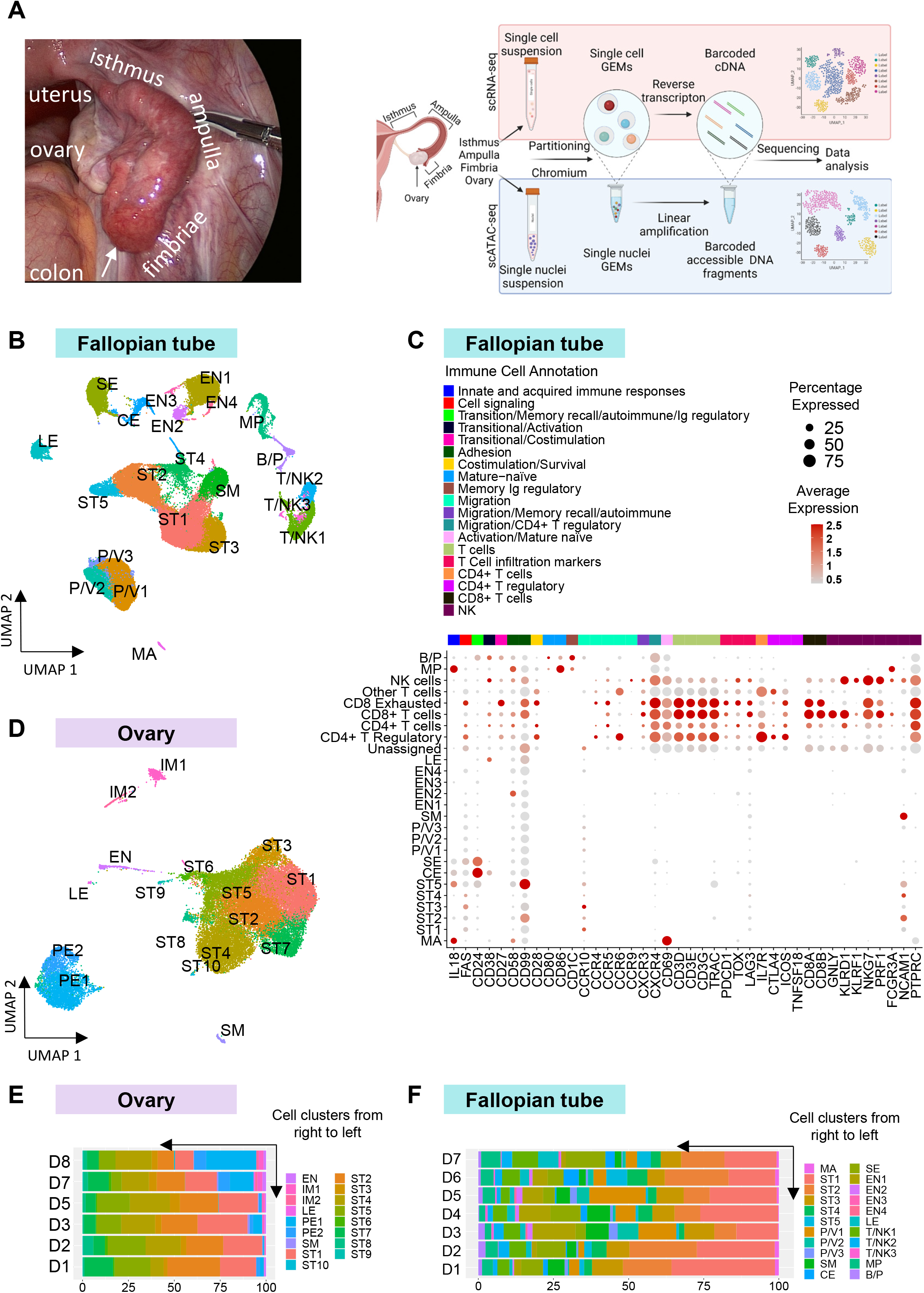
Single-cell RNA sequencing (scRNA-seq) reveals cell types of the normal human postmenopausal fallopian tubes and ovaries. A) Left. Intraoperative image of the ovary, right side of uterine fundus, isthmus – the fallopian tube segment closest to the uterus, ampulla, and the distal end of the fallopian tube (fimbriae). The fimbrial end extends over the ovary allowing direct contact between the epithelium of the fimbriae (arrow) and the ovarian surface. Right. Schematic of the female reproductive tract and experimental workflow for scRNA-seq and scATAC-seq. B) Cell types found in the normal postmenopausal fallopian tube. UMAP plot showing the 22 cell clusters identified in the fallopian tube using scRNA-seq. Data includes all three anatomic regions (fimbriae, ampulla, and isthmus) for seven donors. C) Dot plot showing common gene expression markers found in immune and non-immune cell sub-types in the fallopian tube as identified by HIPPO analysis. D) Cell types found in the normal postmenopausal ovary. UMAP plot showing the 17 cell clusters identified in the ovary using scRNA-seq. Data includes a total of six donors. Cell types are abbreviated as follows: ST1-10 = 10 clusters of stromal cells, SM= smooth muscle cells, EN= endothelial cells, LE= lymphatic endothelial cells, IM1/2= 2 clusters of immune cells. PE1/2 = perivascular endothelial cells. E, F) Relative abundance of 17 cell clusters in the postmenopausal ovary (E) and the 22 cell clusters found in the fallopian tube (F) using scRNA-seq. The graphs show the individual percentage of each cell type by the individual donor. Abbreviations: ST= stromal cells, T/NK= T cells and NK cells, SE= secretory epithelial cells, LE= lymphatic endothelial cells, SM= smooth muscle cells, MP= macrophages, P/V= pericytes and vascular smooth muscle cells, CE= ciliated epithelial cells, EN= endothelial cells, B/P= B cells and plasma B cells, MA= mast cells, NEW1-3= Unknown clusters detected from Cancer cell 2020 dataset only.

A deadly disease threatening the health of postmenopausal women is ovarian cancer^3^. There are more than 30 histological subtypes of benign, borderline, and malignant ovarian tumors listed in the WHO clinical staging system, and many of these have no identified cell of origin. The most common subtype, epithelial high-grade serous ovarian cancer, is now believed to arise from the epithelial cells in the fallopian tube; however, there is still uncertainty since stromal cells may also play a role in the origin of the disease^4,5^. A comprehensive characterization of all cell types in the normal postmenopausal fallopian tube and ovary will form the basis for understanding how these organs undergo malignant transformation.

Single-cell techniques have quickly expanded our understanding of the underlying cellular heterogeneity of ovarian cancer, but there is limited information about the cellular composition of the fallopian tube and ovary. Moreover, most current studies have been limited to single-cell RNA-sequencing (scRNA-seq). While these reports provide critical information, it is becoming increasingly evident that gene expression data alone may not adequately define cell types or elucidate the transcriptional regulations involved in the development, maintenance, and aging of the female reproductive system. Epigenetic information, such as chromatin accessibility is likely to be essential for depicting a more complete regulatory landscape. In this study, we used a combination of technologies, including Drop-seq, 10X single-cell RNA-seq (scRNA-seq), and single-cell assay on transposase accessible chromatin (scATAC-seq) to generate a high-quality cell atlas that consists of four tissue sites from the reproductive tract of postmenopausal women. This atlas, by far the most comprehensive map of the postmenopausal female reproductive system to date, will pave the way to a better understanding of the pathophysiology of many diseases affecting older women.

## Results

### Characterizing canonical cell types in fallopian tube and ovary

This study of the normal postmenopausal fallopian tube and ovary included 8 non-smoking postmenopausal women over 55 years of age undergoing surgery to treat vaginal prolapse without any major macroscopic and histologic abnormality (**Table 1**). Fresh tissues were transported expeditiously from the operating room to the laboratory for cell dissociation and used for scRNA-seq and scATAC-seq. To ascertain changes in cell type, gene expression, and chromatin accessibility over the length of the fallopian tube (approximately 11 cm), tissue samples were taken from the isthmus, which is close to the uterus, the ampulla, which is mid-tube, and the fimbriae, at the end of the tube (**Fig. 1A**).

In total, 18 fallopian tube tissue samples from 7 donors were profiled using scRNA-seq. Among them, 5 tissue samples from 2 donors (D1, D2) were processed using Drop-seq, and the remaining were processed using 10x genomics. After removing doublets and cells with high mitochondrial content, 60,574 cells from all three anatomic sites of the FT were retained for cross-sample integration and downstream analysis. We identified 22 clusters (**Fig. 1B**) across the three anatomical subsections of the FT and all donors, using unsupervised clustering and canonical marker genes (**Fig. S1A, Table 2**). The 22 FT clusters were classified into 11 major cell types (**Fig. S1C**) based on the expression of marker genes (**Table 3**): ciliated epithelial (CE), secretory epithelial (SE), smooth muscle (SM), pericyte/vascular (P/V1-3), endothelial (EN1-4), lymphatic endothelial (LE), stromal (ST1-5), mast (MA), and immune cells: T and natural killer (NK) cells (T/NK1-3), macrophages (MP), B and plasma (B/P) cells. Significant differences in gene expression were seen between the different subgroups of P/V cells. Secretory epithelial (SE) and ciliated epithelial (CE) cells could not be further subclustered using scRNA-seq, which was not the case in studies of FT tissue from premenopausal patients^6,7^. While all ST cells expressed *COL1A1* and *PDGFRA*, 2 subgroups become apparent. ST1/3/4 strongly expressed the mediators of Wnt signaling, POSTN and SFRP4. ST2/5 expressed *DCN, NCAM1, CD99*, and a stem cell marker, *CD34* (**Fig. 1C, Fig. S1A, B**).

The UMAP clustering analysis (**Fig. 1B**) worked well in the definition of major cell types, but failed to resolve finer sub-clusters within some major cell types, such as T and NK cells. We, therefore, applied an alternative approach, HIPPO, which can resolve cellular heterogeneity by iterative feature selection and clustering^8^. While Seurat uses a common set of genes to define all clusters in the dataset, HIPPO applies a hierarchical strategy and represents each new cluster using a different set of markers (**Table 3**) by a zero-inflation test. When HIPPO was applied to T and NK cells for finer sub-clustering, multiple subsets of T cells were revealed, including CD4+ T cells, CD4+ regulatory T cells, and CD8+ T cells (**Fig. 1C**). Two sub-clusters from this analysis can be attributed to CD8+ T cells by the expression of *CD8A* and *CD8B. PDCD1, TOX*, and *LAG3* were expressed in one CD8+ T-cell cluster, but were not expressed or weakly expressed in the other, suggesting that this second cluster might represent a subpopulation of exhausted CD8+ cells^9^. We also noted the expression of *CD99* or *MIC2*, immune-related genes that increase T cell adhesion and apoptosis, in ST2 and ST5 cells, and *CD24*, a cancer stem cell marker, in SE and CE cells.

scRNA-seq was also performed on the ovaries of 6 donors using Drop-seq (D1, D2) and the 10X genomics (D3, D5, D7, D8) assay. After removing doublets and high mitochondria content cells, we obtained 26,134 ovarian cells for downstream analysis. Unsupervised clustering of the cells from the ovaries across donors yielded 17 clusters (**Fig. 1D, Fig. S1D, Table 4**), which were classified into 6 major cell types (**Fig. S1E**) using selected marker genes: stromal (ST1-10), perivascular endothelial (PE1-2), smooth muscle (SM), endothelial (EN), lymphatic endothelial (LE), and immune (IM1-2) cells. Consistent with the high expression of the known stromal marker, decorin (*DCN*), in ST cells in scRNA-seq, most stromal cells in the ovary expressed DCN, as shown using RNA-FISH (**Fig. S1F**). Most cells in the ovary are stromal cells, but there is an unexpectedly large fraction of perivascular endothelial cells (**Fig. 1D**). We did not find evidence of epithelial cells, as the ovary is covered by a single layer of epithelial cells that comes off during surgery.

For both the FT and the ovary, there was some variability in cell-type composition between donors (**Fig. 1E, F**), but in general, they followed the overall cell-type composition seen in the computationally pooled samples. The FT and ovary expressed no oocyte markers (**Fig. S1G**), consistent with the postmenopausal state of the ovaries^10^. There was no appreciable difference in cell types or compositions between Drop-seq and 10x Genomics 3’ RNA-seq data for both organs.

We compared our scRNA-seq dataset from the FT with published data of scRNA-seq in the FT^7,11^. **Fig. S1H** shows the UMAP of datasets of Hu, *et al*.^11^, Dinh, *et al*.^7^, and our FT data. There was minimal overlap in UMAP space with fresh and cultured fallopian tube epithelial cells from Hu, *et al*., possibly due to different single cell sequencing approaches. However, we found significant overlap with the scRNA-seq profile from one postmenopausal donor in the Dinh, *et al*.^7^ study. Comparing our data with the published studies, we see significant overlap in the expression of genes implicated in high-grade serous ovarian cancer (data not shown).

### Characterization and gene expression in different anatomic regions of the human postmenopausal fallopian tube

Driven by known structural and functional differences in anatomic regions, we further characterized the three anatomic regions: the isthmus, the ampulla, and the fimbriae of the FT (**Fig. 1A, 2A**), at the cellular and molecular levels (**Fig. 2B, C**). Overall, each major cell type was present in roughly similar proportions in all three regions of the FT (**Fig. 2A, B**). **Fig. 2C** shows the normalized expression levels of key genes in the major cell types of the isthmus/ampulla/fimbriae and the percentage of cells in the respective clusters expressing them. Secretory and ciliated epithelial cells shared several pan-epithelial cell markers, *EPCAM, KRT8/18/19, FOLR1, SLP1, WFDC2* in all anatomic regions^12^. Secretory epithelial cell specific markers present in all three anatomic regions, that were not expressed in ciliated cells, included *KRT7, OVGP1*, and *MSLN. RASGEF1B* is strongly expressed in the isthmus but reduced in the ampulla and absent in secretory epithelial cells of the fimbriae. The complement gene, *C3*, and the tumor suppressor gene, *CSMD1, were* also reduced in secretory epithelial cells of the fimbria (**Fig. 2C**). Expression of markers specific to ciliated epithelial cells included known markers ^7^ (*CAPS, FOXJ1*) and new markers (*PIFO, TMEM190, SNTN*). The histone-related gene, *HIST1H4C*, is expressed in ciliated epithelial cells of the isthmus but not in the ampulla and the fimbria, and *RTN1* is not expressed in ciliated epithelial cells of the fimbriae (**Fig. 2C**).

**Figure 2:**
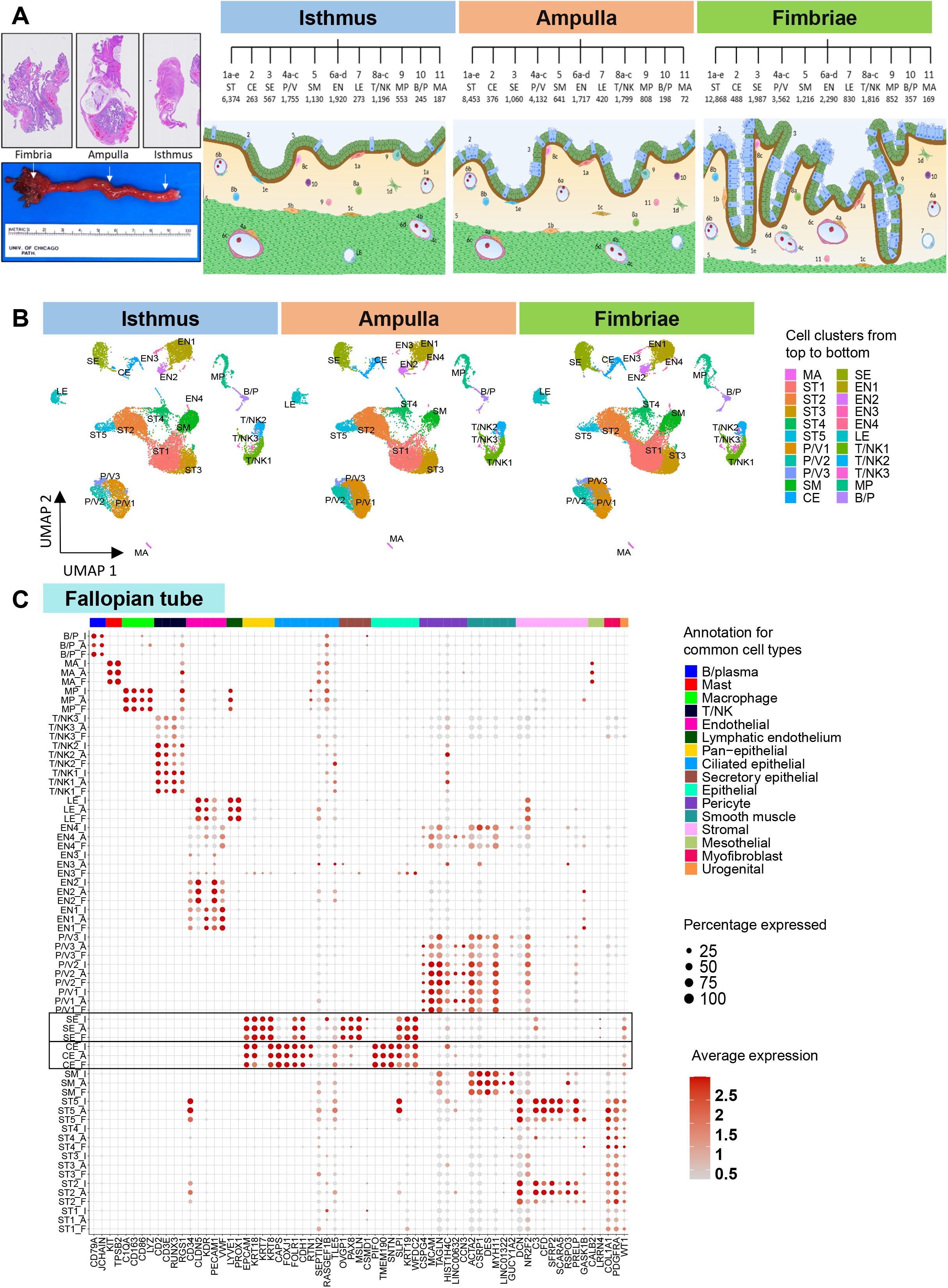
Gene expression in the isthmus, ampulla, and fimbrial regions of the postmenopausal fallopian tube. A) Left panel: Gross anatomic image of a normal fallopian tube indicating the anatomic regions sampled and their corresponding cross-section H&E (1:40). Right panel: Graphical depiction of cell sub-clusters and number of cells identified in each sub-cluster by anatomic region (isthmus, ampulla, and fimbria), using scRNA-seq. B) UMAP of the 22 cellular clusters identified by scRNA-seq divided by anatomic sites in the fallopian tube (isthmus, ampulla, and fimbriae). C) Dot plot of normalized expression levels of marker genes in major cell types in the isthmus (I), ampulla (A), and fimbria (F) and cell types expressing them. Secretory (SE) and ciliated epithelial (CE) cells are framed. Abbreviations of cell types are described in Figure 1.

Though the three regions (**Fig. 1A, 2A**) essentially share a similar pattern of major cell types, we noticed FT heterogeneity in non-epithelial cell subtypes, including T/NK3 and P/V3, ST2, and ST5. All stromal cells in the fallopian tube (ST1-ST5) were characterized by *COL1A1* and *PDFGRA* expression. Within the stroma cluster, the ST2 and ST5 subcluster expressed *DCN, which* is important for collagen assembly and as an inhibitor of angiogenesis and tumorigenesis, through all three anatomic regions. ST5 cells in the isthmus and ampulla express *SLP1, CD34, C3, CFD, SFRP2*, and *SCARA5*, while these are lost in the fimbriae; they only express GASK1B. The gene expression pattern of pericytes, smooth muscle, and endothelial cells was mostly unchanged across the isthmus, ampulla, and fimbriae (**Fig. 2C**).

Finally, we performed immunohistochemical staining in the isthmus/ampulla/fimbriae and ovary of our patients to verify the presence of select cell types identified by scRNA-seq (**Fig. S2A**). The staining confirmed the canonical gene expression for each cell type and anatomic region on the protein level. Vimentin was expressed in stromal and epithelial cells. EPCAM was expressed primarily in the ampulla and fimbriae epithelial cells, and only minimallly expressed in the isthmus. We also show PAX8 protein expression in secretory epithelial cells and FOXJ1 protein expression in ciliated epithelial cells in all 3 regions of the FT. CD45 and low levels of CD68 expression were seen in some cells in all regions of the FT and the ovary (**Fig. S2A**).

### Correlation of scRNA-seq results with genome-wide association studies (GWAS)

Gene expression measurements at single-cell resolution in the female reproductive system provide unique opportunities to pinpoint gynecological disease associations with specific cells^13^. We examined 83 genes (**Table 5**) related to disease-causing variants, identified from GWAS, of 8 gynecological diseases. These diseases, which included several carcinomas and endometriosis, substantially affect postmenopausal women, and manifest and likely originate in the FT or ovary. We included endometriosis because it sometimes persists into menopause and can be a precursor for endometrioid and clear cell ovarian cancer^14^. We found 65 of 83 risk-associated genes expressed in at least one cell type in the FT and 64 of 83 in the ovary (**Fig. 3A**). Not surprisingly, most risk-associated genes only manifest high expression in 1 or 2 cell types, and their expression patterns vary in the FT and ovary. Normal ciliated and secretory epithelial cells expressed several serous high-grade ovarian cancer-related genes^11^ including keratins (*KRT17, KRT23*), metabolism (*ALDH1A1, ALDH3B2)*, immune (*HLA-DQA1, HLA-DPA1*), and stemness related genes (*LGR5, CD44*) (**Fig. S3B**). Surprisingly, only *MSI2*, a stemness gene expressed in high-grade ovarian cancer, showed expression in both the secretory and ciliated cells of the FT (**Fig. S3A**). Moreover, we discovered several new relationships of high-grade serous ovarian cancer associated genes and benign cell types in the FT: Ciliated epithelial cells in all three anatomic regions of the FT highly expressed *STK33, ADGB*, and *CCDC170* (**Fig. 3A**). These genes were not represented in secretory epithelial cells, which are currently thought to be the cell of origin for high-grade serous ovarian cancer, suggesting a contribution from ciliated epithelial cells to the pathogenesis of the disease^15^.

**Figure 3:**
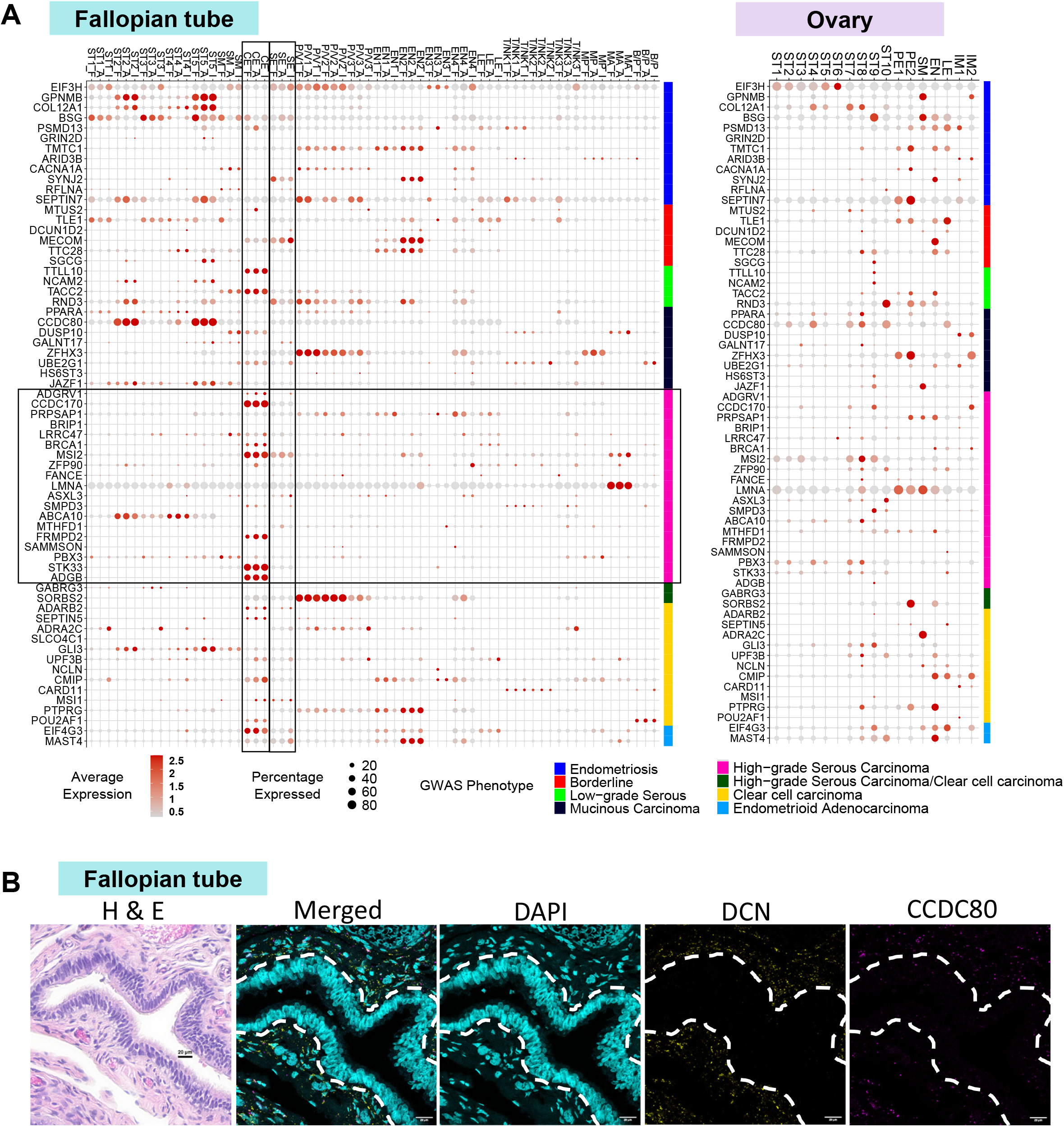
scRNA expression of genes identified in genome-wide association studies (GWAS) in different regions of the postmenopausal fallopian tube and the ovary. A) Dot plot showing cell type-specific expression of disease-specific genes as curated by GWAS in the ovary and fallopian tube by anatomical site (isthmus, I; ampulla, A; fimbriae, F). B) Hematoxylin and eosin (H&E) and RNA-fluorescent in situ hybridization (FISH) of decorin (DCN) and CCDC80 in the fallopian tube. Every dot corresponds to one RNA transcript. Nuclei are stained with DAPI (cyan). Transcripts for DCN are shown in yellow and CCDC80 in magenta. Dashed white lines separate the epithelium from stroma in the fallopian tube (scale bars 20 μm). Abbreviations of cell types as described in Figure 1.

The mucinous carcinoma-associated gene, *CCDC80*, a known tumor suppressor in ovarian cancer^16,17^ and expressed in a subset of stromal cells (ST2, ST5) in the FT (**Fig. 3A**) was confirmed using FISH (**Fig. 3B**). Endometriosis associated genes, *COL12A1, GPNMB, BSG* and *SEPTIN7*, all show moderate to high expression in stromal cells in the FT (**Fig. 3A**), though in different subsets. They also manifest high expression in various non-immune, non-epithelial cell types in the ovary: *COL12A1* in several stromal compartments, *GPNMB* and *BSG* in smooth muscle cells, and *SEPTIN7* in perivascular epithelial cells (**Fig. 3A**). This is particularly interesting because endometriosis is thought to arise from displaced endometrial cells and not ovarian stromal cells^18^.

In addition to cell-type and tissue-site-specific differences in the ovary and FT, there were significant differences in the expression patterns of GWAS genes between individual donors. Most major cell types had a unique expression signature (**Fig. S3A, B**). Expression patterns within most cell sub-clusters are consistent, in that the same set of GWAS genes are expressed in each woman, albeit to various levels (e.g., FT: P/V1/2/3, T/NK1/2/3, ST1/3, ST2/5; ovary: ST1/2/3/4/5/6/7/8, PV1/2, IM1/2). There were, however, a few notable exceptions, in which at least one woman expressed a distinct set of GWAS genes (FT: EN1/2, ST4; ovary: ST9/10, EN, LE). Donors D3 and D5 expressed a relatively higher number of ovarian cancer –associated genes in the FT. In the ovary (**Fig. S3A**), ST9 was the subcluster expressing the most ovarian cancer-associated genes (**Fig. 3A**). These analyses yield an expression map of risk genes in a cell type-specific and tissue site-specific fashion, providing hypotheses for the cell of origin of gynecological disease.

The ovary and the fallopian tube are hormone-responsive organs, but the current clinical thinking is that, once a woman is a few years into menopause, they are “non-functional”. To determine if the postmenopausal FT and ovary express hormone receptors that circulating hormones can potentially activate, we systematically analyzed the expression of 63 receptors (**Table 6**) in the different cells of the FT and ovary and found that 60 and 59 receptors, respectively, are expressed by at least one cell type (**Fig. S3C**). The expression of several receptors involved in metabolic regulation of FT cells (e.g., the adiponectin, insulin, and gastro inhibitory polypeptide receptors) suggests that FT cells are metabolically active and have the receptors to respond to systemic hormonal changes. Ciliated epithelial cells express the adiponectin, estrogen, insulin, and oxytocin receptors. All stromal cells in the FT express progesterone receptors (PGR, PGRMC1) and the androgen receptor, while these receptors are absent in the ovary (only ST9 has some PGR expression). The estrogen receptor was expressed in all stromal cells of the FT, but not in the stromal cells in the ovary (**Fig. S3C**). qRT-PCR of primary stromal cells (FTSCs) and epithelial cells (FTECs) from the fallopian tube and ovarian stromal cells (OVST) showed high expression of PGR in OVST and high ESR1 expression in FTEC and FTSC (**Fig. S3E**). Though few cells were stained for estrogen and progesterone receptors in the ovaries, as is expected in post menopause, it was intriguing to find strong immunohistochemical staining for estrogen and progesterone receptors in epithelial and stromal cells of the isthmus/ampulla/fimbriae (**Fig. S3D**).

### Ligand-receptor interactions between different cell types in fallopian tube and ovary

To understand the interactions between different cell populations and how they jointly create the FT microenvironment, we inferred ligand_receptor interactions across all cell types in various anatomic sites using the scRNA-seq as input for the CellPhoneDB software package^19^ (**Fig. 4, Table 7**). In the FT, the most robust interactions were observed between ST5, secretory epithelial, and endothelial cells across all anatomic regions (**Fig. S4A**). Most strong interactions between stroma and endothelial cells, as well as ciliated and secretory epithelial cells, across all anatomic sites, were driven by CD74, a chaperone receptor. CD74 regulates antigen presentation for immune responses that can bind to MIF, a cytokine inhibiting immune function by increasing the prevalence of a highly immune-suppressive population of myeloid-derived suppressor cells^20^ (**Fig. 4A**). We discovered significant differences when comparing ligand-receptor interactions in the isthmus and ampulla with those in the fimbriae. In the fimbriae, ciliated cells secreted COPA, a protein involved in endocytosis that could potentially bind to the CD74 receptor on ciliated, endothelial, and stromal cells. In contrast, interactions between MIF and TNF receptor family members detected in the isthmus, and ampulla were absent in the fimbriae (**Fig. 4A**).

**Figure 4:**
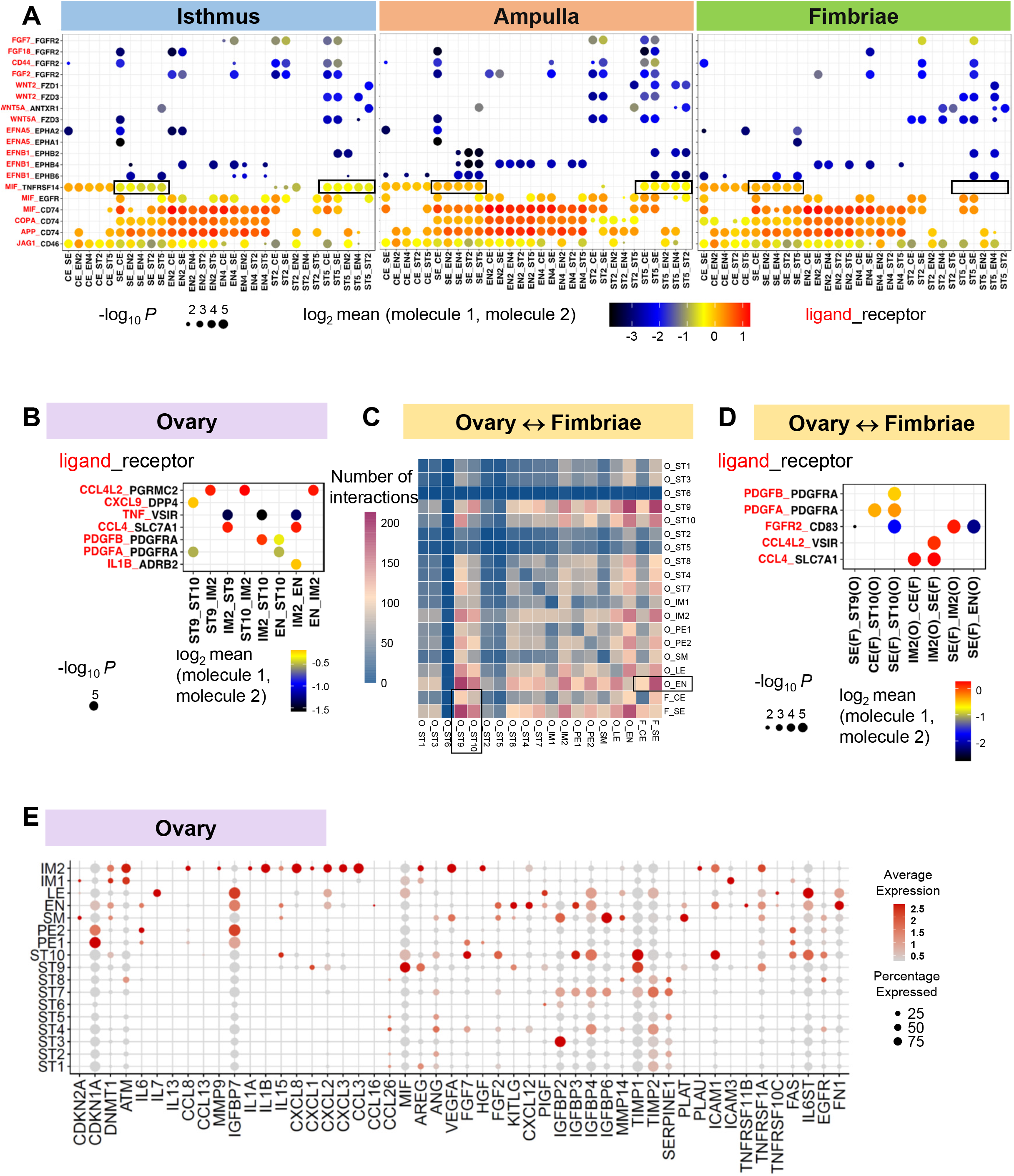
Ligand-receptor interactions and senescence-related gene expression in the fallopian tube and ovary. A, B) Ligand: receptor interactions between different cell populations in the postmenopausal fallopian tube (A) and ovary (B) as detected by CellphoneDB. Y-axis: Ligands (red), receptors (black). X-axis: Cell types with ligand-receptor interactions separated by underline. Black boxes indicate examples of significant changes between anatomic sites. C) Ovary – Fallopian tube interactions. Heatmap showing the number of interactions detected by CellPhoneDB among different cell types in the ovary and with SE and CE cells in the fimbria of the fallopian tube. D) Ligand-receptor interactions between fimbriae and ovary, as detected by CellPhoneDB. E) Aging and senescence-related gene expression in the ovary. Abbreviations of cell types as described in Figure 1.

In the ovary, we found fewer receptor | ligand interactions. Of note is the interaction between stromal and endothelial cells secreting cytokines (*CCL4, CCL4L2*) that bind to immune cells (**Fig. 4B**). We show the total number of imputed ligand-receptor interactions in donors D3 and D5 in isthmus/ampulla/fimbriae, and ovary (**Fig. S4B**) where interesting anatomical region and patient-specific differences are evident. In all anatomical regions of the FT in donor D3, the SE, ST5, EN2, and EN4 cells show the largest number of interactions among themselves and with other cells (P/V3, T/NK3), while donor D5 showed the highest level of interactions in the isthmus only. In the ovary, D3 and D5 show few cellular interactions, consistent with their postmenopausal state (**Fig. S4B**).

Because the ovary and fimbriae are in direct contact (**Fig. 1A**), we determined the interactions between epithelial cells (SE, CE) in the fallopian tube and the different cell types in the ovary. We found potentially strong interactions between ovarian ST9/10 and endothelial cells with both secretory and ciliated epithelial cells in the FT (**Fig. 4C, S4C**). These interactions were driven by receptor-ligand pairs that play important roles in the epithelial-to-mesenchymal transition, including *PDGFR, FGFR*, and *CCL4* (**Fig. 4D**). CCL4, potentially secreted by immune cells, binds to the SLC7A1 receptor expressed on ciliated and secretory epithelial cells. SLC7A1 is important for glucose and amino acid transport across the plasma membrane (**Fig. 4D**).

The women participating in this study represent an aging population (55 years and above), which is why we examined genomic signatures related to aging and senescence in our samples (**Table 5**). Senescent cells can cause tissue damage by secreting high inflammatory cytokines and growth factors as part of the senescence-associated secretory phenotype (SASP)^21^. We see expression of several SASP-associated genes, such as *SERPINE1, TIMP1, TIMP2, IGFBP2/3/4* in FT and ovarian stromal cells (many clusters), and *VEGFA, FGF7* and *EGFR* in FT stroma (**Fig. 4E, S4D**). We also note the expression of several *CCL* and *CXCL* genes in immune cells from the FT (MP) and ovary (IM2), potentially undoing some of the deleterious effects of senescence (**Fig. 4E, S4D**). As with the scRNA-seq results, we saw significant variability between different donors in both FT and ovary (**Fig. S4B, C**).

### Chromatin accessibility analysis of the postmenopausal fallopian tube and ovary

Single-cell ATAC-seq (10X Genomics) was performed on a subset of donors to match their scRNA-seq data. Cells harvested from FT and ovary were split for scRNA-seq, and the remaining underwent nuclei preparation and transposition treatment for scATAC-seq. The cells from isthmus/ampulla/fimbriae from five women (D3, D4, D5, D6 and D8) were processed separately and integrated *in silico*. A total of 41,515 cells passed our QC criteria for scATAC-seq^22^. For each individual sample, we integrated scATAC-seq and scRNA-seq, using label transfer^23-25^. When both data types were available for any given donor, we used the cluster information obtained from scRNA-seq as a reference and explicitly searched for the best-matched cluster for every single cell in scATAC-seq. In total, we identified 40,803 cells from the FT (**Fig. 5A**) from scATAC-seq with matched scRNA-seq compartments. **Table 8** shows label transfers between scRNA-seq and scATAC-seq standalone clusters, the majority of scRNA-seq and scATAC-seq clusters matched well, confirming the high quality of both data sets. Then Accessibility matrices constructed at gene levels were integrated across samples and batch effects were corrected using *Harmony*^26^ and piped into the downstream analysis. We identified 25 clusters by unsupervised clustering, which could be further classified into the same 11 major cell types seen in scRNA-seq analysis (**Fig. S1C**) based on chromatin accessibility (scATAC-seq) matched to cell type labels (scRNA-Seq) (**Fig. 5A**). Similarly, we performed scATAC-seq on ovaries obtained from three donors (D3, D5, and D8). The same QC, integration, and clustering procedures used for FT yielded 18,335 cells that could be grouped into 13 cell clusters that belong to five major cell types: stromal, perivascular, endothelial, smooth muscle, and immune cells (**Fig. 5B**). In contrast to the ovarian scRNA-seq data (**Fig. S1D**) we did not identify lymphatic endothelial cells.

**Figure 5:**
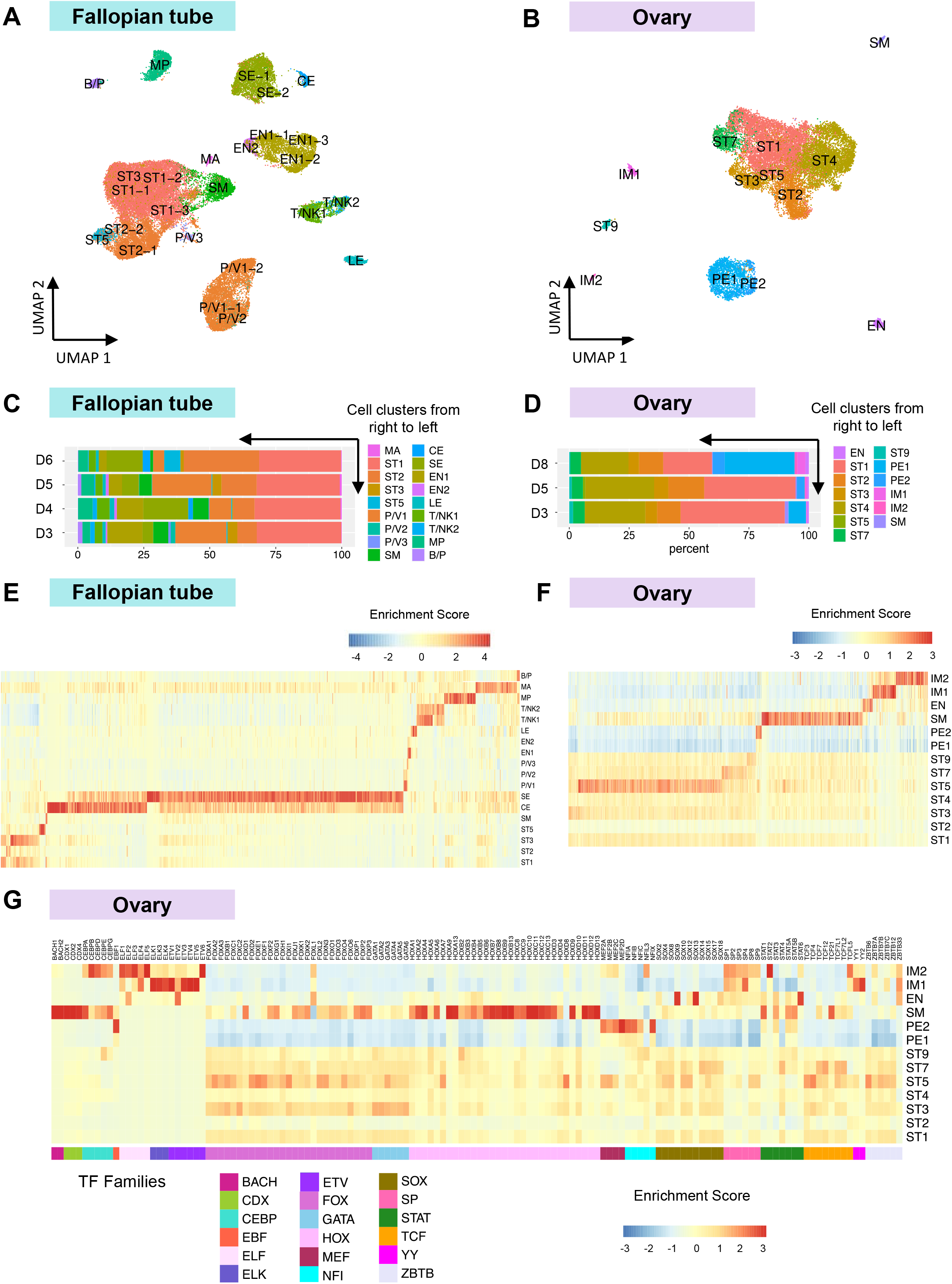
Single-cell Assay of Transposase-Accessible Chromatin sequencing (scATAC-seq). Annotation of cell-types by label transfer from scRNA-seq in the postmenopausal fallopian tube and ovary. A, B) UMAP plot profiling of (A) 40,803 cells from 4 donors identifying 18 major cell clusters in the fallopian tube (isthmus, ampulla, fimbria), and (B) 18,335 cells from 3 donors identifying 13 major cell clusters in the ovary using scATAC-seq. The labels before “-” are transferred from scRNA-seq data and scATAC-seq specific cluster labels are added as subgroup numbers after “-” (for example, there are now secretory epithelial (SE) cell clusters -1 and -2). C, D) Relative abundance of the 18 cell clusters in the postmenopausal fallopian tube and 13 cell clusters found in the ovary using scATAC-seq after integrated analysis (label transfer from scRNA-seq). The graph shows the individual percentage of each cell type by the donor. E, F) Heatmaps depicting transcription factor activity by cell type in the fallopian tube (E) and the ovary (F). The heatmaps show all 869 motifs available in the cisBP database. G) Heatmap depicting specific transcription factors in scATAC-seq data from the postmenopausal ovary from the cisBP database. Abbreviations of cell types as described in Figure 1.

We identified sub-clusters for several cell types in the FT that could not be differentiated by gene expression but were separated by measuring chromatin accessibility. For example, in the FT, EN1 could be further resolved into three sub-clusters EN1-1, EN1-2 and EN1-3, Secretory epithelial cells could be resolved into sub-clusters SE-1 and SE-2, and P/V1 into P/V1-1 and P/V1-2 by scATAC-seq (**Fig. 5A**). P/V3 are in close proximity to stromal cells, consistent with their similar gene expression^27^. EN3/4 and T/NK3 in the FT and ST6/8/10 and lymphatic epithelial cell types in the ovary that were identified by scRNA-seq (**Fig. 1B, D**) could not be detected by scATAC-seq. This could be due to the similarity of these cell types regarding chromatin accessibility, differences in the number of cells profiled, or inherent limitations of the method, as the percentage of cells for these subtypes were small in scRNA-seq (**Figs. 1E, F**). The percentages of identified subtypes in scATAC-seq also vary among donors (**Fig. 5C, D**).

To fully characterize the regulatory landscape in the FT and the ovary, we estimated the activities of 870 transcription factors (TF) listed in the *cisBP* database^28^ in a cell-type-specific fashion (**Fig. 5E, Table 9**). In the ovary, we found ST5, SM, PE2 and immune cells to be the most transcriptionally active cell types (**Fig. 5F**). The ELK, ELF and SP1 TF families are enriched in endothelial and immune cells, and the EBF1 and MEF families are enriched in PE1 and PE2 (**Fig. 5G**). Most of the ST cells in the ovary show low TF activity except for ST5, which shows relatively high activity in the GATA, FOX and TCF families. The ZBTB and YY families are enriched in immune cells, which is consistent with ZBTB7B and ZBTB7A’s role in regulating the development and/or differentiation of conventional CD4/CD8 αβ+ T cells and the role YY1 plays in regulating broad general processes throughout all stages of B-cell differentiation^29,30^.

FOXL2 is essential for embryogenesis, cell differentiation, and tumorigenesis. The highly conserved nature of this gene, and its limited expression, predominantly in the ovary, suggests that it is a key factor throughout ovarian development. Our data shows that most FOX TF family members ^31^ show high accessibility in the ovary, particularly for FOXL2 in ST5 (**Fig. 5G**). At the same time, SOX9 acessibility is repressed in stromal cells in the ovary (**Fig. 5G**). We also detect high *FOXL2* expression (↑) and *SOX9* supression (↓) in several stromal clusters in FT (ST1/3/4) (**Fig. S5A**). This pattern is reversed in secretory and ciliated epithelial cells (↓ *FOXL2*, ↑ *SOX9*) of the FT (**Fig. S5A**).

Ablation of FOXL2 in the adult mouse ovary causes immediate induction of the transcription factor, SOX9, leading to ovary-to-testis trans-differentiation. We observe a similar pattern of high *FOXL2* and low *SOX9* expression in several ST clusters of the FT when reviewing the browser track plots of FOXL2 and SOX9 accessibility and expression (from scATAC-seq and scRNA-seq data, respectively) for each cell type and anatomic location (**Fig. S5B**). In SE and CE of the FT, *SOX9* shows a strong chromatin accessibility signal, while *FOXL2* shows no signal. *FOXL2* chromatin accessibility is found in ST2/5 in the fimbriae but is absent in the isthmus and ampulla. Yet, *FOXL2* is expressed across isthmus/ampulla/fimbriae for ST1/3/4. This is yet another instance of expression patterns differing in ST2/5 compared to ST1/3/4 in the FT as a whole (**Fig. S1B**) and by anatomic location (**Fig. S5B**).

### scATAC-sequencing allows the identification of cell-type and location-specific regulatory elements

The chromatin landscape in the fallopian tube displays similar patterns in the isthmus, ampulla, and fimbriae (**Fig. 6A**). However, the TF motif analysis revealed that, while regulation patterns across cell types are almost identical in the isthmus and ampulla, they are strikingly different in the fimbriae (**Fig. 6B, C**). For example, SE-1 and SE-2 show much greater chromatin accessibility in the isthmus and ampulla than in the fimbriae. Mast cells are more actively regulated in the fimbriae compared to those in the isthmus and ampulla (**Fig. 6B**). In contrast, immune cells, including B/P, T/NK, and MP, exhibit similar patterns across all 3 anatomic regions, suggesting that the entire FT may respond to immune stimuli similarly (**Fig. 6B**). Equally important, other cell types in the isthmus and ampulla (SM and EN2) and the ST1-3 cells across all FT appear inaccessible (**Fig. 6B, C**). Surprisingly, both ciliated and secretory epithelial cells show lower TF activity of SOX, STAT, and ZBTB TF in the fimbriae, while FOX families are unchanged and are high in ciliated epithelial cells in all 3 anatomic regions of the FT. In respect to endothelial cells, we observed an increased chromatin accessibility signal in SOX family members from isthmus/ampulla to fimbriae. The EN1-1/2 subclusters are characterized by their differences in accessibility of pro-inflammatory STAT transcription factors, which play a role in endothelial cell dysfunction during aging^32^. EN1-1 and EN1-2 both deriving from the same EN1 cluster in scRNA-seq (**Fig. 1B**) via label transfer, must necessarily have similar gene expression. Using scATAC-seq, we found that, in fimbriae, the EN1-2 subclusters have high EFL2 and TCF7L1 activity, while EN1-1 has high YY1 and YY2 activity. These findings suggest that endothelial cells express similar genes but are regulated by different TFs (**Fig. S6A**). For ETV2, TF activity progressively decreases from isthmus to fimbriae in EN1-3 and increases from isthmus to fimbriae in EN1-1 (**Fig. S6A**).

**Figure 6:**
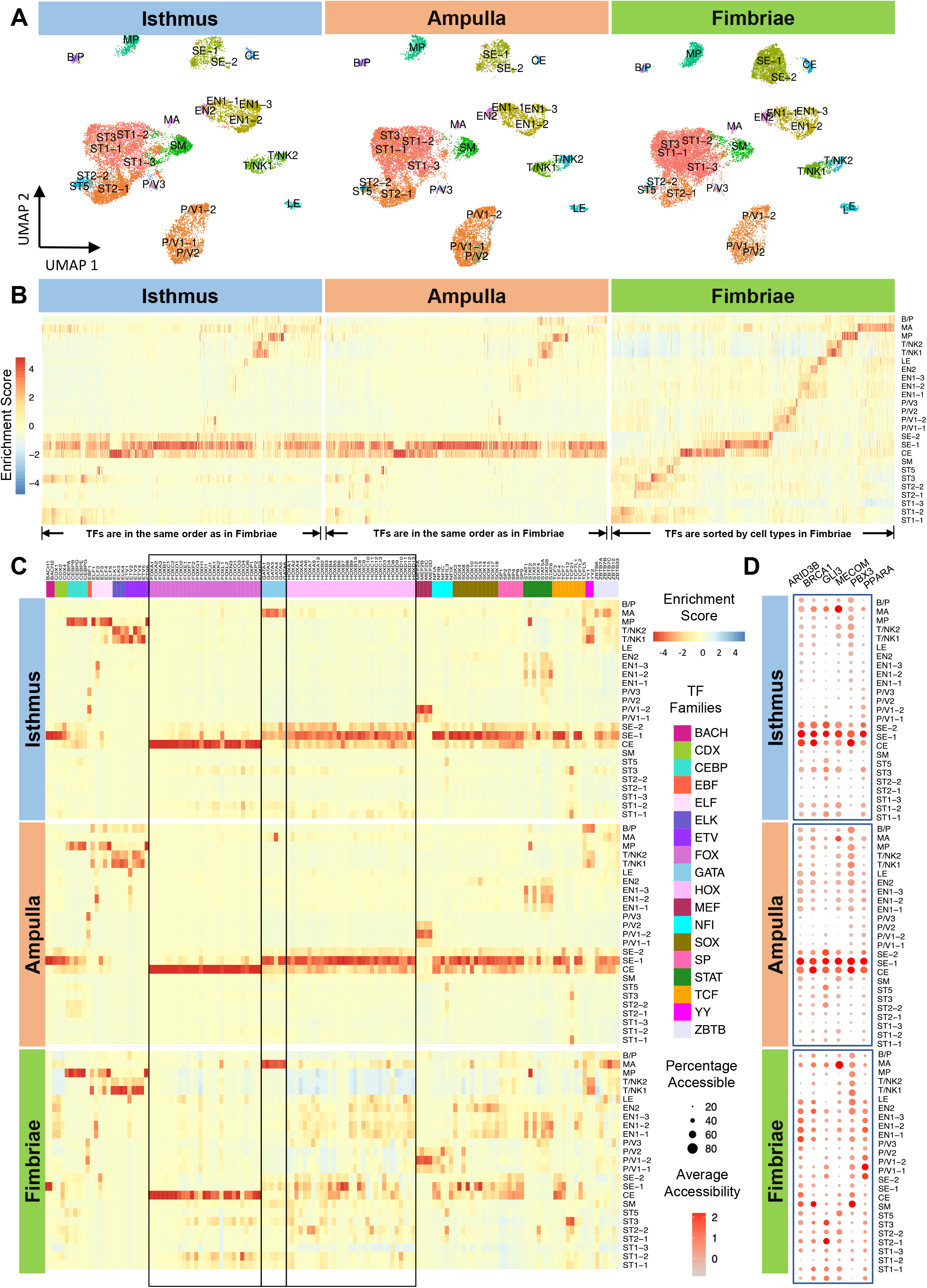
Clustering, cell-type annotation, and transcription factor (TF) analysis of scATAC-seq in the different anatomical regions of the postmenopausal fallopian tube. A) UMAP plot showing the 18 major cell clusters identified in the isthmus, ampulla, and fimbria by scATAC-seq after label transfer from scRNA-seq. B) Heatmap showing TF activity in the isthmus, ampulla, and fimbriae by cell type. Enrichment results were obtained from the cisBP database and contain 869 transcription factor motifs. The order of TF is the same across anatomical sites. C) Heatmap showing selected transcription factors in the isthmus, ampulla, and fimbria. D) Dot plot showing chromatin accessibility of TF of GWAS genes in Figure 3 by anatomical site and cell type. Abbreviations of cell types as described in Figure 1.

Known EMT-related TFs, including BACH1 (**Fig. 6C**), SNAI1, SNAI2, ZEB1, and TWIST1 (**Fig. S6B**), show enriched activities in SE, but not in stromal cells, suggesting EMT related TF activity specifically in FT epithelial cells. The JUN/FOS family of transcription factors show enriched activities in ST5 and SE (**Fig. S6B**) where different isoforms of JUN and FOS proteins are activated in epithelial (SE-1) and stromal cells (ST5). SE-1 expressed FOS/JUND, which has known inhibitory transcriptonal function^33,34^. The TF RUNX3, which plays a critical role in the differentiation of premenopausal FT^7^, was not expressed in CS and SE, but showed increased chromatin accessibility in T/NK cells.

We then interrogated the activities of TFs in our curated list of GWAS genes (**Table 5**), including ARID3B, BRCA1, GLI3, MECOM, PBX3, and PPARA. GWAS-related TF accessibility is similar in the isthmus and ampulla but is strikingly different in the fimbriae. In the isthmus and ampulla, most TF show high activity in CE and SE, but are less active in the other cell types. In contrast, in the fimbriae, TF accessibility is more robust in most non-epithelial cells (EN, ST2/3/5) (**Fig. 6D**). In summary, there are several cell types showing major differences in chromatin accessibility in the fimbriae (macrophages ↑, endothelial ↑, CE ↓, SE ↓). These differences were not evident when analyzing TFs using scRNA-seq data, and only became evident when inspecting cluster-specific TF activities identified from scATAC-seq data.

## Discussion

The fallopian tube and ovary, which are crucial to the female reproductive system, are thought to have little physiologic function after menopause. However, these organs are often the origin of diseases that either persist through menopause (e.g., endometriosis) or newly develop (benign or malignant tumors) later in life. Without an appreciation of the healthy state of an organ, it is difficult to understand any pathological condition, and therapies are most effective if they address the underlying changes from the normal to the diseased state.

With this in mind, we set out to systematically characterize the gene expression and regulatory landscape of all cell types in the human postmenopausal FT and ovary at single-cell resolution, to develop a resource that could be a starting point for all scientists studying female well-being and disease. The different cell types we identified in the FT overlapped almost entirely with a recent report on one postmenopausal FT^7^. Because the histologic appearances of the FT close to the uterus and at the fimbrial end are so different, we further examined several anatomic regions of the FT separately. Interestingly, within all 11 cell types, there was little change in gene expression between the three anatomic areas of the FT. However, there were remarkable differences in chromatin accessibility between the fimbriated end of the FT when compared to the isthmus and ampulla. Several cell types in the fimbriae showed strong enhancer activity that included macrophage, endothelial, perivascular, and some stromal cell types. Endothelial cells in the fimbriae showed increased accessibility to the SOX transcription factor (TF) family, known to activate endothelium^35^. Since the SOX TF family is involved in tissue repair/regeneration and endothelial mesenchymal transition, upregulation in fimbriae is probably a reaction to the chronic exposure of the fimbriae to the peritoneal cavity fluid or to the aging of the FT. Interestingly, ciliated and secretory epithelial cell types that were previously implicated in disease etiology showed limited chromatin accessibility to STAT, and ZBTB family of TFs in the fimbriae, indicating resting/senescent epithelium^36^. Senescent cells can cause tissue damage by secreting high levels of inflammatory cytokines and growth factors as part of the senescence-associated secretory phenotype (SASP)^21^. SASP-associated genes, such as *VEGFA, FGF7*, and *EGFR* are expressed in FT stroma, and others, such as *SERPINE1, TIMP1, TIMP2, IGFBP2/3/4* are expressed in both FT and ovarian stromal cells (many clusters).

In contrast, immune cells throughout all anatomic regions of the FT, as well as the ovary, remain transcriptionally active and express antigens and cytokines, indicating that both organs are not immunologically inert. We also noted the expression of several *CCL* and *CXCL* genes in immune cells from the FT (macrophages) and ovary, potentially undoing some of the deleterious effects of senescence and aging. We did not find any of the subclusters reported by Ulrich et al in their elegant study of premenopausal fallopian tubes^6^. Nor did we find the subclusters detected by Dinh, *et al*. in the premenopausal ovary^7^. Using scRNA-seq, we found just one type of ciliated and secretory epithelial cells each; but with scATAC-seq, we could resolve SE cells into SE-1 with accessibility by multiple FOS/JUN heterodimers and EMT regulating TF, while SE-2 cells had low occupancy for most of these TF. With menopause, the fallopian tube epithelium loses the sophisticated sub-differentiation of epithelial cells required to perform its reproductive functions.

For our study, we used fresh tissue from healthy postmenopausal women that was apparently normal on microscopic examination and expressed clinically established immunohistochemical markers. We learned that the concept of “normal” tissue is relative; while all cell types were represented in every patient, there was wide variation in gene expression between patients, consistent with reports on “normal” lung^13^ and kidney^23^ tissue. Even when all cells of a “normal” cell type had the same gene expression, we found that subclusters varied in their transcriptional regulatory patterns. For example, scATAC-seq allowed the identification of three endothelial subclusters (EN1-1/2/3) in the FT, that were each regulated by a different set of transcription factors but showed similar gene expression. This observation could explain the remarkable adaptability of “normal” cells to stressors in the local microenvironment, highlighting the startling self-healing potential of utilizing redundant regulatory pathways. Our data suggest that the integration of scRNA-seq and scATAC-seq is essential for deciphering the cellular complexity of tissues, which would not be possible with either of the assays alone. A limitation of our integration was our inability to confirm gene expression levels corresponding to TF activity for cellular subtypes identified from scATAC-seq alone. This limitation can be overcome in future studies by using recent multi-omics assays that measure gene expression and chromatin accessibility simultaneously from the same cell.

When correlating our scRNA-seq results in the FT with endometriosis^37^ associated risk genes, we identified *GPNB*, a gene implicated in the inflammatory response, in stromal subclusters ST2/5, and *BSG*, an immunoglobulin family member frequently detected in ectopic endometrial tissue, in ST1/3/5^38^. *BSG*, also known as EMMPRIN or CD147, promotes cell proliferation by blocking apoptosis and upregulating MAPK signaling in human endometrial epithelial cells^39^. In the ovary, *EIF3H*, a translation initiation factor implicated in endometriosis, was expressed in 6 stromal subtypes. It is intriguing that most of the candidate-risk genes found in endometriosis are expressed in stromal cells. We will have to await more data on the pre-menopausal FT and ovary, as well as the results of functional studies to fully understand the significance of these findings.

Several ovarian cancer risk genes implicated by GWAS studies were also expressed in stromal cell populations in the FT. Stromal clusters ST2/5 expressed *CCDC80*^16^, which is part of a tumor microenvironment gene signature derived from the ovarian cancer related TCGA data^40^. Surprisingly, most GWAS-derived risk genes in the normal postmenopausal FT were not expressed in secretory cells, which were suspected of being the ovarian cancer cell of origin^41^. Instead, they were found in the ciliated epithelial cells (*TTLL10, MECOM, TACC2, CCDC170, MSI2, BRCA1, STK33, ADGB, CMIP*).

We also present the first scRNA-seq and ATAC-seq atlas of the human postmenopausal ovary. Most cells in postmenopausal ovary are stromal cells, but there are also a substantial number of endothelial cells, consistent with the very rich blood supply to the ovary from the aorta and the uterine artery. There are few immune cells in the postmenopausal ovary, and, unlike the ovaries of aged peri-menopausal non-human primates^42^, no follicles were detected, suggesting complete follicular atresia in human menopause. The human postmenopausal ovary had a much higher variation in stromal gene expression than that of the monkeys, but in contrast to peri-menopausal monkey ovary very few genes were associated with senescence or DNA oxidation. In general, the stromal cells in the postmenopausal ovary are not very transcriptionally active; however, the ST5 cluster retains marked accessibility for all FOX and GATA TF family members compared to the other stromal sub-clusters. Based on the ATAC-seq data, endothelial, smooth muscle, and immune cells in the postmenopausal ovary have high chromatin accessibility, so by no means should the ovary be considered a quiescent organ, especially given the strong expression of several hormone receptors potentially bound by circulating hormones (e.g. leptin, androgen, prostaglandin receptors etc.). It is fascinating that, in the postmenopausal setting, ovarian cells have almost no estrogen receptor expression, while all FT stromal cells have high estrogen and progesterone receptor expression. Unopposed estrogen production during menopause (e.g. from adipose tissue) might cause mutagenic effects in the tube, as described for the postmenopausal endometrium in which estrogen contributes to the transformation of the resting epithelium^43^. The fimbrial end of the FT and the ovary are next to each other and depend on close anatomic and functional interactions to transfer the oocyte during ovulation from the ovary to the FT. Our results indicate that the FT epithelial cell types (CE, SE) and ovarian stromal and immune cells may interact directly through complementary ligand-receptor expression. These findings provide the basis for follow-up studies using high-content spatial transcriptomic/ proteomic imaging.

With this study, we focused on the postmenopausal FT and ovary and were able to characterize the largest number of FT and ovarian cells in a single integrated study to date. We have characterized and integrated the regulatory and transcriptional landscape of the postmenopausal FT and ovary at single-cell resolution, providing an important reference dataset to study reproductive physiology and disease. The challenge that will remain is the integration and translation of this information into functional studies *in vivo* to ultimately provide effective therapies for previously intractable disease states.

## Acknowledgments

We thank Dr. Mark Eckert for his incredible help and dedication during the initial phase of this project. We thank Gail Isenberg for her help with editing the article.

Grant funding: This work was mainly supported by the Chan Zuckerberg Initiative (to E.L., O.B., M.W. M.E., M.C.). M.C. is supported by the NIH (R01 GM126553 and R01 HG011883), the National Science Foundation (NSF 2016307), and the Sloan Research Fellowship Program. A.B. is supported by NIH DP2AI158157. E.L. is supported by R35CA264619, RO1CA211916, RO1CA237029, and the Ovarian Cancer Research Alliance.

We thank the Genomics Facility, Human Tissue Resource Center and Pritzker Nanofabrication Facility at the University of Chicago for equipment and services, and Dr. Ran Zhou for help with single cell experiments. Microfluidic devices were fabricated in the University of Chicago. Patients from The University of Chicago OB/GYN department donated tissue for this study.

## Author contributions

E.L., A.B., and M.C. jointly oversaw project design and analysis. Y.L., L.Z., and M.C. performed all computational analyses. H.E., M.J., S.A., M.E., J.X. and S.O. performed experiments including tissue dissociation, Drop-seq, and 10X Genomics single-cell RNA and ATAC-seq assays. M.W. performed the FISH experiments. R.L. and A.J.B. read histology and immunohistochemistry. D.G. and S.I. consented and performed the surgeries on the women donating the tissue. E.L., A.B., M.W. and M.C. wrote the manuscript. All authors reviewed and agreed on the final version of the submitted manuscript.

## Methods

### Materials

Five μm sections of formalin-fixed paraffin-embedded tissues from postmenopausal women were stained with hematoxylin and eosin or with commercially available antibodies using the Leica Bond RX automated stainer (Leica Biosystems) and reviewed by gynecologic pathologists. The following antibodies were used for immunohistochemistry: FOXJ1 (clone 2A5, eBioscience, 1:500), CD68 (clone PG-M1, Dako, 1:200), CD45 (clone 2B11+ PD7/26, Dako, 1:100), pan cytokeratin (clone AE1/AE3, Biocare medical, 1:200), vimentin (clone V9, Dako, 1:2000), PR (clone 16, Novocastra, 1:300), ER (clone 6F11, Novocastra, 1:180), PAX8 (10336-1-AP, Proteintech, 1:1000), EPCAM (HPA026761, Prestige antibodies, 1:75), WT1 (clone 6F-H2, Invitrogen, 1:400), CK7 (clone OV-TL12/30, Dako, 1:1000).

### Tissue acquisition and dissociation

Fallopian tube and ovary samples were collected from female patients who underwent elective surgical hysterectomy for benign indications (vaginal prolapse, incontinence) and were having these tissues removed as part of their normal surgical procedure performed at The University of Chicago Medical Center (**Table 1**). Tissue was collected from postmenopausal women defined as absence of a menstrual cycle for greater than two years and/or absent menses with accompanying symptoms of menopause. Signed informed consent was obtained from the patient prior to the elective procedure. All procedures involving the human samples were conducted in accordance with the Institutional Review Board at the University of Chicago.

We followed stringent, standardized criteria for sample collection and began dissociating tissues within 20 minutes after surgical removal. Tissue specimens from eight patients were collected and processed (**Table 1**). The tissue was removed surgically and processed immediately after the tissue was amputated from its surrounding tissue and blood supply^44^. The three anatomic regions (I, A, F) were further dissected and dissociated independently into single cell suspensions for higher anatomical resolution, to obtain scRNA-seq and scATAC-seq data of high quality. Due to the variability in tissue mass and cell viability, we were able to obtain sufficient number of cells from only two of the three regions of the FT from certain donors (**Table 1**). We also note that in cases where the ovaries were not removed, we obtained tissues from the fallopian tubes without matched ovarian cells.

100 mg of ovarian tissue and 2-3 mm cross-sections of each fallopian tube segment (100 mg each of isthmus, ampulla, and fimbriae) were digested independently^45^. Each tissue slice was rinsed with a fetal-bovine serum enriched DMEM (SIGMA) to remove blood and mucus and surrounding parametrial tissue was trimmed. The dissociation was a two-stage protocol separating each tissue into epithelial and fibroblast components with an initial epithelial digestion with pronase at 37° for 30 minutes^46^. Epithelial cells were filtered out and the remaining stroma-fibroblast supernatant underwent second digestion with tissue DNAse, collagenase IV, hyaluronidase digestion at 37° for 30 minutes. At the end of the digestion, the epithelial and stromal-fibroblast components were combined and passed through 70 μm filter. The cell suspension was spun at 400 rcf for 7 minutes and the cell pellet was re-suspended in DMEM+FBS. To remove remaining red blood cells from processing, the red blood cell lysis solution (EasySep RBC Depletion Reagent, Stemcell Tech #18170) was used. The cells were imaged and cell counts were obtained. During dissociation, we aimed for cell viability over 85% (assessed by Trypan blue, Sigma T8154 and/or Annexin V, BD 556547 staining) and cells of diverse morphologies (assessed by bright-field imaging) to meet QC criteria.

### RNA scope – Fluorescent in situ hybridization

RNAscope was performed on 5 μm sections of formalin-fixed and paraffin-embedded tissues using the Leica Biosystems’ BOND RX System and ACD biotech user manual (Document number: 322800-USM) for the RNAscope LS multiplex fluorescent reagent kit user manual for BDZ 11. The following probes were purchased from ACD biotech for RNAscope: DCN (Cat.: 589528) and CCDC80 (Custom probe targeting 2600-3557 of NM_199511.3 in C3). Images were acquired using a Nikon Eclipse Ti2 microscope and processed using the Nikon software NIS-Elements (version: AR5.30.05 64-bit).

### Single-cell RNA-sequencing

scRNA-seq was performed on 8 donors using Drop-seq (2 donors-D1, D2) and 10X genomics (6 donors, D3 - D8 using the 10x Genomics 3’ RNA-seq assay).

#### Drop-seq experiment

Drop-seq experiments were performed as previously described^47^. Briefly, cells and oligonucleotide barcode beads were loaded at concentrations of 100,000 cells/ml in PBS-BSA and 120,000 beads/ml in Drop-seq lysis buffer in 3 ml syringes. Droplets were generated using a 125-micron microfluidic device at 16 ml/hr (oil), 4 ml/hr (cells) and 4 ml/hr (barcode beads, MACOSKO-2011-10 (V+), ChemGenes Corp.) with ∼15 minutes per collection. Following collection, drops were broken and barcoded beads with mRNA hybridized onto them were collected and washed. Barcoded cDNA attached to the beads or STAMPs were generated by reverse transcription, treated with Exonuclease I and the number of STAMPs was counted. 5000 STAMPs were aliquoted per well in a 96-well plate and the cDNA attached to the STAMPS were amplified through 15 PCR cycles. Supernatants from each well were pooled and cleaned with Ampure beads. Purified cDNA was quantified using Qubit 3.0 (Invitrogen) and 450-650 pg of each sample was used as input for Nextera reactions (12 cycles). Tagmented libraries were quantified using Agilent BioAnalyzer High sensitivity chip before submission for sequencing on Illumina’s NextSeq 500, using 75 cycle v3 kits. Paired end sequencing was performed with 20 bp for Read 1 and 60-64 bp for Read2 using a custom Read1 primer, GCCTGTCCGCGGAAGCAGTGGTATCAACGCAGAGTAC and 5% Illumina PhiX Control v3.

#### 10X Genomics 3’ scRNA-seq experiment

Single-cell suspensions were transported to the lab in warmed media to preserve viability. The cells were washed once with PBS + 0.4% BSA and resuspended in PBS + 0.4% BSA to achieve a target cell count of 700-1200 cells/μl. An appropriate amount was loaded, based on target cell counts, according to the Chromium Next GEM protocols. Single-cell suspensions from each sample were loaded onto the fluidic chip at X cells/uL were constructed into barcoded 3’ scRNA-seq libraries using the Chromium Next GEM kit, v3 targeting 8,000 cells/sample, except for two samples from the isthmus (D4 and D5) that had low number of cells where only 2000 cells were targeted. Sequencing libraries were constructed following 10X genomics protocols with 15 amplification cycles. Bioanalyzer 2100 (Agilent) traces were used to evaluate cDNA and final sequencing libraries. The libraries were sequenced through the University of Chicago core facility using a PE75 run on the Illumina NextSeq 500 or NovaSeq platforms. Libraries were sequenced at 30,000-50,000 reads per cell according to the manufacturer’s recommendations.

### scRNA-seq data analysis

The original raw BCL (binary base call) sequencing data were converted and demultiplexed into FASTQ files by a wrapper function, *cellranger mkfastq*, from Cell Ranger software^48^ developed by 10x genomics (https://support.10xgenomics.com/single-cell-gene-expression/software/overview/welcome). Then the raw sequencing reads were aligned to the human reference genome hg38 and then filtered and quantified as UMI counts using barcode information via *cellranger count*. We further applied the following QC criteria to the UMI count matrix using an in-house pipeline: 1) we require cells expressing at least 200 gene features and each gene feature present in at least 3 cells; 2) we remove doublets and triplets identified by *DoubletDecon* ^49^; 3) cells with ≥20% mitochondrial contents were filtered out as poor-quality cells with low viability. Each UMI count matrix, with cells from a certain anatomical site of a donor, is log-normalized using a procedure^50^ implemented with *Seurat*^51^ function *NormalizeData* and *FindVariableFeatures*. Cells from different samples were integrated using *Seurat* function *FindIntegrationAnchors*^52^, where top 2,000 highly variable gene features expressed across cells were used as anchors for pairwise sample integration. Prior to dimensionality reduction, a linear transformation scaling procedure was applied to remove unwanted variations. We performed dimensionality reduction using both PCA and UMAP. Shared nearest-neighbor (SNN) graph was constructed for graph-based clustering via *Seurat* function *FindNeighbors* and *FindClusters* correspondingly, at resolution of 0.5 for a relatively dense clustering. Cell clusters obtained from unsupervised learning (top 100 differentially expressed genes in the FT and ovary are listed in **Tables 2, 4**) were further manually annotated using canonical markers of different cell types curated from the literature (**Table 3**). Sub-clustering of immune NK/T cells was performed by HIPPO^8^, a method solving cellular heterogeneity using zero proportions instead of gene variance. The raw UMI counts of selected cells were input to HIPPO for sub-clustering analysis, with the number of clusters specified 2 times higher than the number of clusters originally identified from Seurat. To avoid early stop before the specified number of clusters, we used a z-score threshold of 1 with default outlier proportion of 0.001%. Obtained sub-clusters are further characterized using a set of additional immune markers (**Table 3**).

Our scRNA-seq data was compared with datasets of normal epithelial cells (Hu, *et al*., Cancer cell 2020) (https://www.ncbi.nlm.nih.gov/geo/query/acc.cgi?acc=GSE139079^11^) and one postmenopausal patient from Dinh, *et al*., Cell Reports, 2021 (https://www.ncbi.nlm.nih.gov/geo/query/acc.cgi?acc=GSE151214)^7^.

### GWAS analysis

We curated a list of 84 genes that contained disease-causal variants for 8 disease categories from public GWAS databases. We checked the expression levels of these genes in our scRNA-seq data. Among these, 6 genes (GPC5, CALHM3, PA2G4P2, SNTG1, KRT18P55, PKD1L1) were barely expressed in all cell types and were removed from further exploration. This resulted in a total of 78 GWAS genes (**Table 5**) for downstream comparisons and display.

### Ligand-receptor analysis using CellPhoneDB

The normalized gene expression data were used as the input for CellPhoneDB. The ligand-receptor interaction analysis utilizes CellPhoneDB function curated by CellPhoneDB database v2.0.0. Significant (P < 0.05) receptor - ligand interactions between any two cell types in different anatomy sites in fallopian tube were displayed (**Table 7**). Similar analyses were conducted and displayed for interactions of cell types in ovary, and interactions between CE and SE in fimbriae and cell types in ovary^19^.

### Single cell-ATAC-seq

#### Experiment

Fresh single-cell suspensions were lysed on ice for 4 minutes to obtain intact nuclei. The nuclei were tagmented at 37 °C for 1 hr according to standard protocol for the Chromium Next GEM kits (10X Genomics) to generate scATAC-seq libraries with 6,000-8,000 nuclei per sample. All downstream procedures were performed following standard manufacturer’s protocols. Agilent 2100 Bioanalyzer traces were used to evaluate final library quality. The libraries were sequenced following 10X Genomics guidelines on Illumina NextSeq 500 and NovaSeq platforms through the University of Chicago Core Facility. Libraries were sequenced at 25,000-30,000 read pairs per nucleus according to the manufacturer’s recommendations.

#### Pre-processing and quality control^22^ of scATAC-seq data

Raw sequencing data from each sample were aligned to human hg38 reference genome using *cellranger ATAC 1*.*2*. R Seurat v4.0^51,53^ and *Signac v1*.*4*.*0*^53^ were used for further analysis. High quality cells, defined as cells with peak region fragments > 3000, peak region fragments < 20000, % of reads in peaks > 15, blacklist ratio < 0.05, nucleosome signal < 5 and TSS enrichment > 2, were retained for normalization using term-frequency inverse-document-frequency (TFIDF). The Seurat objects from each sample (after label transfer from scRNA-Seq) were merged based on the common peak set which was created by merging peaks from all the datasets. Dimensional reduction was performed via singular value decomposition (SVD) of the TFIDF matrix and UMAP. Batch effects across samples were corrected by *Harmony*^26^ using *RunHarmony* function on the first 30 latent semantic indexing (LSI) components, excluding the first one because it was highly correlated with the sequencing depth. Finally, gene activity scores were estimated using *Seurat* function *GeneActivity*^51^. Data corrected by Harmony were used for unsupervised clustering analysis using *FindNeighbor and FindClusters* functions in *Seurat*^51^.

#### Integration of scRNA-seq and scATAC-seq datasets

To obtain cell types of scATAC-seq data, cells from the matched scRNA-seq analysis were used as a reference dataset to predict cell types in the scATAC-seq. This prediction used the variable features of the scRNA-seq data as reference, and the gene activity matrix generated using Seurat’s *GeneActivity*^*51*^ from scATAC-Seq data as the query data. Transfer anchors were learned using *FindTransferAnchors*^51^ and cell type labels were predicted using *TransferData*^51^ with the scATAC-seq LSI reduction as *weight*.*reduction* input. Specifically, we assign each cell in the scATAC-seq with a cell type (subcluster) identity from the matching scRNA-seq data based on the first 30 LSI components corrected by Harmony, excluding the first one. Only cells with the prediction score, denoted by *prediction*.*score*.*max* that quantifies the uncertainty with predicted annotations, larger than 0.5 were kept for further analysis. Cell clusters transferred from scRNA-seq were further separated into sub-clusters if supported by unsupervised clustering analysis using scATAC-seq data alone. The label transfer procedure was performed for each individual patient separately (**Table 8**).

#### Transcription factor motif analysis

Transcription factor activities (**Table 9**) were estimated from Harmony integrated scATAC-seq data using chromVAR v3.14^54^. TFs and their binding motifs listed in human_pwms_v2(cisBP)^28^ database was used as another input to chromVAR for positional weight matrix calculation. RunChromVAR ^54^ in Signac was applied to calculate the cell type-specific TF activities and differential activities among cell types were computed with *FindMarkers* with Bonferroni-adjusted p-values < 0.05^23,24^. The total number of motifs in cisBP database is 870, while there are only 869 motifs detectable in fimbriae and since we ordered the heatmaps according to fimbriae, there are 869 motifs shown in heatmaps. To obtain a more comprehensive profiling for JUN/FOS family motifs we also checked JASPAR database^55^, which in total contains 633 motifs, within which 29 are JUN/FOS family motifs, significantly more than the collection in cisBP (8 out of 870). These JUN/FOS motifs have different splice forms resulting in different binding specificity, which were handled as binding variants and could be seen from their names (e.g., FOS::JUN(MA0099.3) and FOS::JUN(var.2) (MA1126.1)). The TF enrichment analysis of all motifs was performed for each cell type in each tissue site separately. The results were then compared and displayed.

## Supplemental tables

**Table 1: Patient cohort and clinical-pathologic information**.

**Table 2: Top 100 differentially expressed genes from all 22 cell clusters identified in the fallopian tube from seven donors (scRNA-seq)**.

**Table 3: Canonical marker genes used to annotate cell types in the fallopian tube and ovary (scRNA-seq)**. The different tabs are for fallopian tube markers, ovary markers and immune and non-immune markers used in HIPPO analysis.

**Table 4: Top 100 differentially expressed genes from each 17 cell clusters identified in the ovary using ovaries from six donors (scRNA-seq)**.

**Table 5: Genome Wide Association Study (GWAS) & aging related genes**. The different tabs are for GWAS genes identified in gynecological diseases & aging and senescence related genes.

**Table 6: Hormone receptor expression and annotation based on scRNA-seq**.

**Table 7: Cellphone DB summary based on scRNA-seq**. The different tabs are for interactions in isthmus, ampulla, fimbriae, ovary and between fimbriae and ovary, respectively.

**Table 8: Label transfers between scRNA-seq and scATAC-seq data**. The label transfer was performed and summarized for each individual donor separately.

**Table 9: Transcription Factor enrichment summary (scATAC-seq)**. The different tabs are for isthmus, ampulla, fimbriae, and ovary, respectively.

