## Supplementary Information for "A molecular atlas of the human postmenopausal fallopian tube and ovary from single-cell RNA and ATAC sequencing"

Figure S1

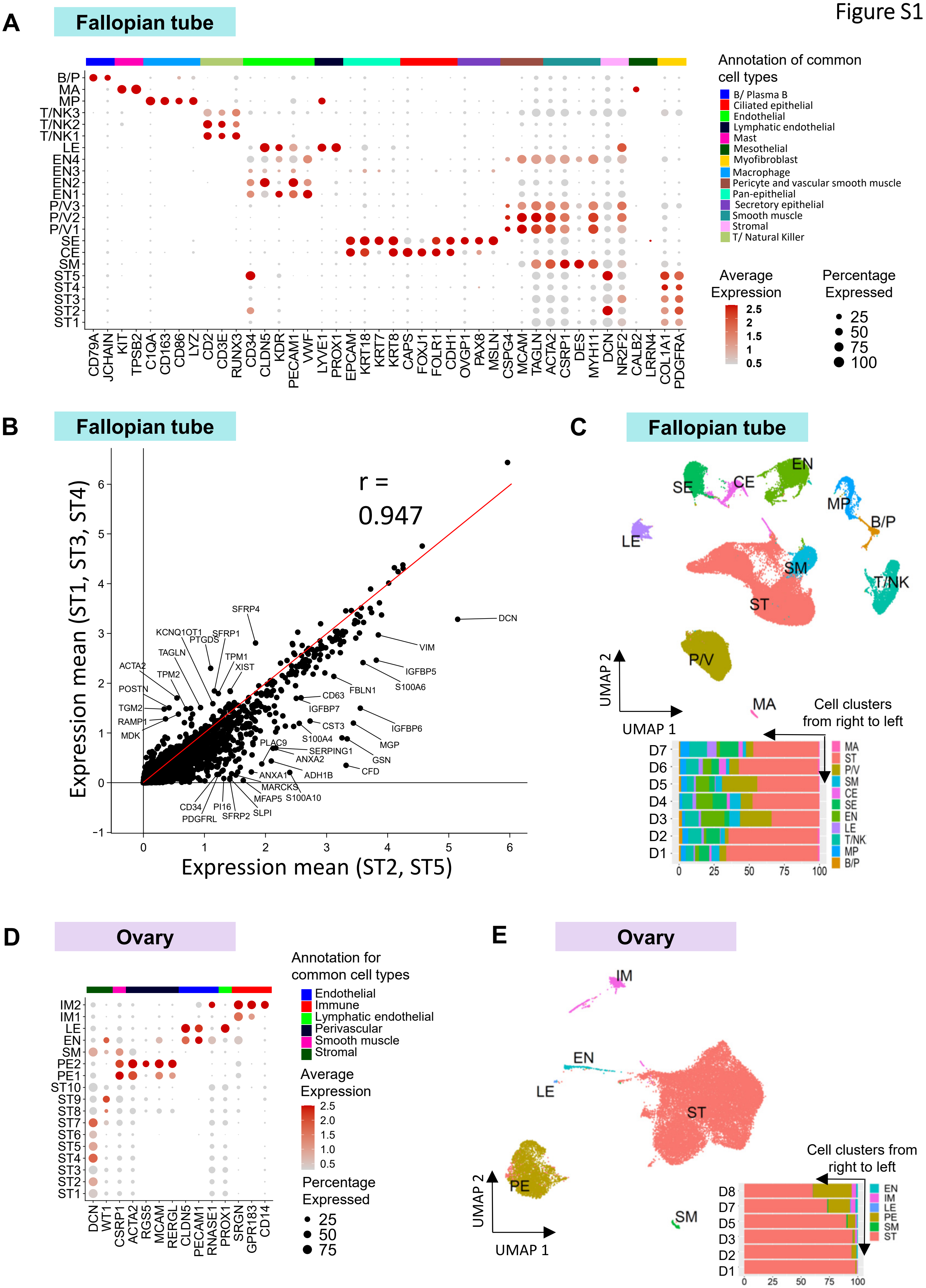

A
Fallopian tube
B/P
MA
MP
T/NK3
T/NK2
T/NK1
LE
EN4
EN3
EN2
EN1
P/V3
P/V2
P/V1
SE
CE
SM
ST5
ST4
ST3
ST2
ST1
CD79A JCHAIN KIT TPSB2 C1QA CD163 CD86 LYZ CD2 CD3E RUNX3 CD34 CLDN5 KDR PECAM1 VWF LYVE1 PROX1 EPCAM KRT18 KRT7 KRT8 CAPS FOXJ1 FOLR1 CDH1 OVGP1 PAX8 MSLN CSPG4 MCAM TAGLN ACTA2 CSRP1 DES MYH11 DCN NR2F2 CALB2 LRRN4 COL1A1 PDGFRA
Annotation of common cell types
B/ Plasma B
Ciliated epithelial
Endothelial
Lymphatic endothelial
Mast
Mesothelial
Myofibroblast
Macrophage
Pericyte and vascular smooth muscle
Pan-epithelial
Secretory epithelial
Smooth muscle
Stromal
T/ Natural Killer
Average Expression
2.5
2
1.5
1
0.5
Percentage Expressed
25
50
75
100
B
Fallopian tube
r = 0.947
Expression mean (ST1, ST3, ST4)
Expression mean (ST2, ST5)
SFRP4
KCNQ1OT1
PTGDS
SFRP1
TAGLN
TPM1
XIST
ACTA2
TPM2
POSTN
TGM2
RAMP1
MDK
DCN
VIM
IGFBP5
S100A6
FBLN1
CD63
IGFBP7
CST3
IGFBP6
S100A4
MGP
PLAC9
SERPING1
ANXA2
GSN
CFD
ANXA1
ADH1B
MARCKS
MFAP5
S100A10
CD34
PI16
PDGFRL
SFRP2
SLPI
C
Fallopian tube
SE
CE
EN
MP
B/P
LE
SM
T/NK
ST
P/V
MA
UMAP 2
UMAP 1
Cell clusters from right to left
D7
D6
D5
D4
D3
D2
D1
0
25
50
75
100
MA
ST
P/V
SM
CE
SE
EN
LE
T/NK
MP
B/P
D
Ovary
IM2
IM1
LE
EN
SM
PE2
PE1
ST10
ST9
ST8
ST7
ST6
ST5
ST4
ST3
ST2
ST1
DCN WT1 CSRP1 ACTA2 RGS5 MCAM RERGL CLDN5 PECAM1 RNASE1 PROX1 SRGN GPR183 CD14
Annotation for common cell types
Endothelial
Immune
Lymphatic endothelial
Perivascular
Smooth muscle
Stromal
Average Expression
2.5
2.0
1.5
1.0
0.5
Percentage Expressed
25
50
75
E
Ovary
IM
EN
LE
ST
PE
SM
UMAP 2
UMAP 1
Cell clusters from right to left
D8
D7
D5
D3
D2
D1
0
25
50
75
100
EN
IM
LE
PE
SM
ST

Figure S1

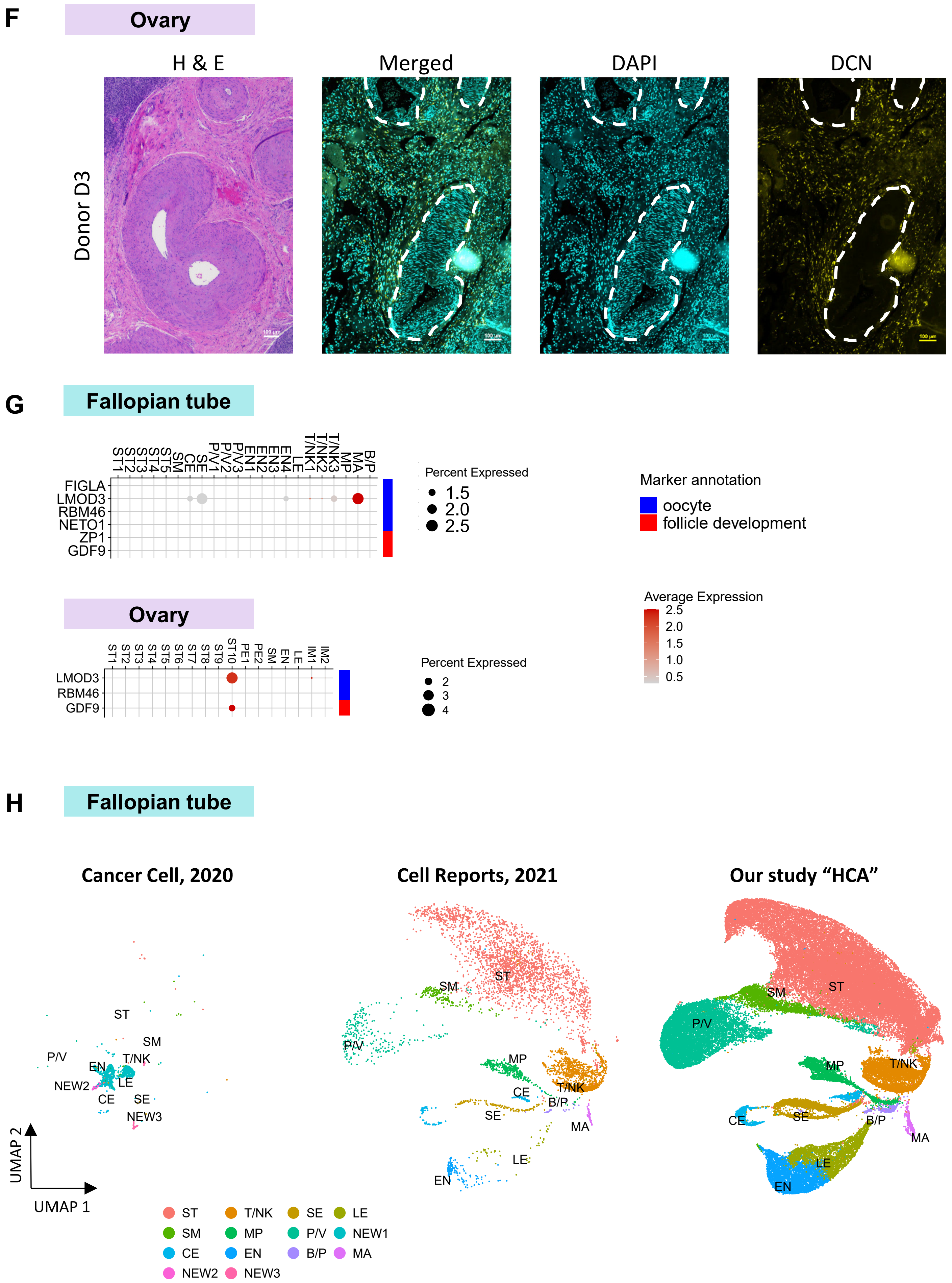

F
Ovary
H & E
Merged
DAPI
DCN
Donor D3
G
Fallopian tube
ST1 ST2 ST3 ST4 ST5 SM CE SE P/V1 P/V2 P/V3 EN1 EN2 EN3 EN4 LE T/NK1 T/NK2 T/NK3 MP MA B/P
FIGLA
LMOD3
RBM46
NETO1
ZP1
GDF9
Percent Expressed
1.5
2.0
2.5
Marker annotation
oocyte
follicle development
Ovary
ST1 ST2 ST3 ST4 ST5 ST6 ST7 ST8 ST9 ST10 PE1 PE2 SM EN LE IM1 IM2
LMOD3
RBM46
GDF9
Percent Expressed
2
3
4
Average Expression
2.5
2.0
1.5
1.0
0.5
H
Fallopian tube
Cancer Cell, 2020
Cell Reports, 2021
Our study "HCA"
ST
SM
P/V
EN
T/NK
NEW2
LE
CE
SE
NEW3
MP
B/P
MA
UMAP 2
UMAP 1
ST T/NK SE LE
SM MP P/V NEW1
CE EN B/P MA
NEW2 NEW3

**A**

Figure S2

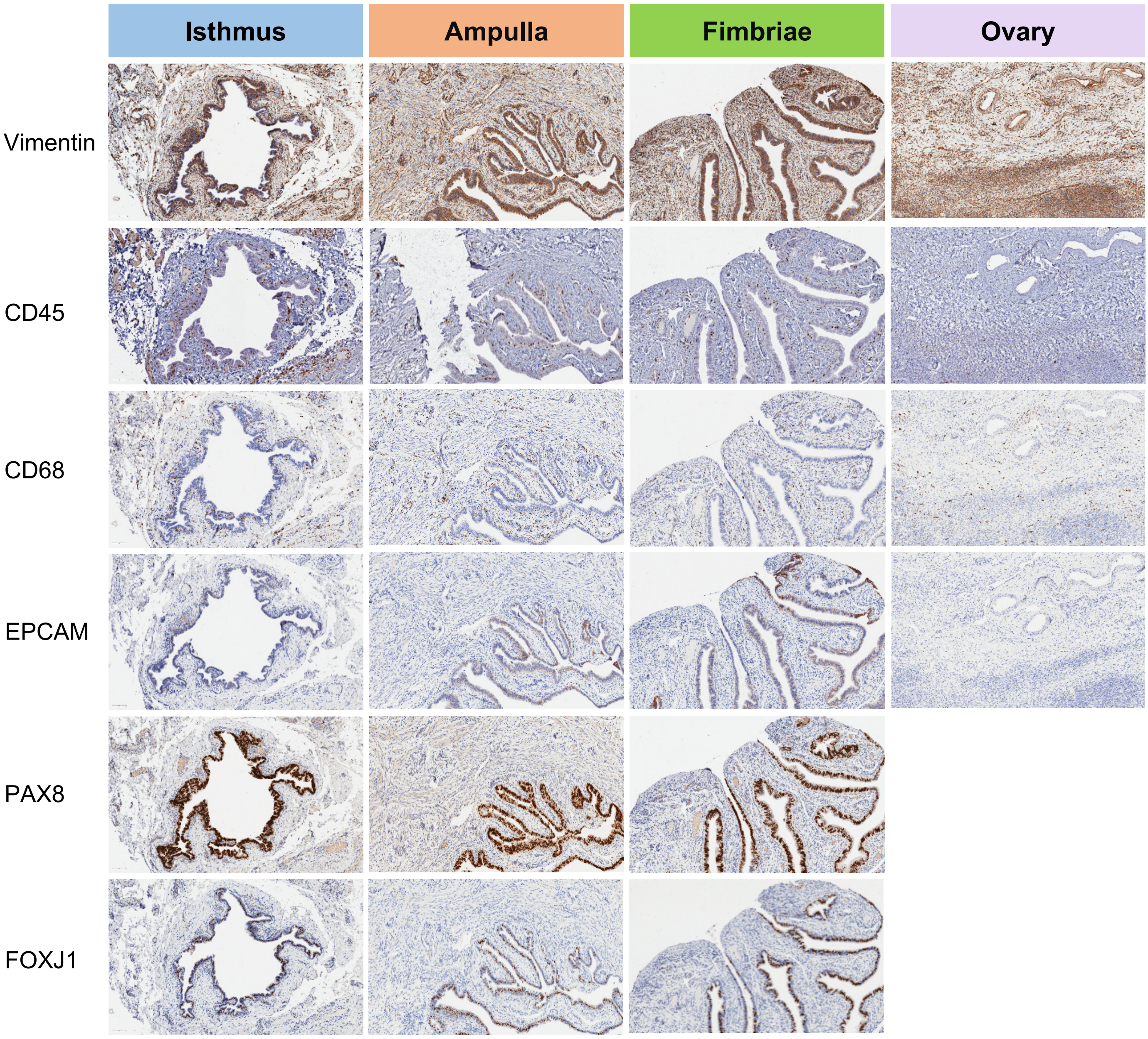


| Antibody/ Target | Description | Cell type |
|---|---|---|
| VIM | An intermediate filament protein that is part of the cytoskeleton. Responsible for maintaining cell shape and integrity of the cytoplasm. | Mesenchymal cell marker |
| EPCAM | Homotypic calcium-dependent cell adhesion molecule. Expressed on normal epithelial cells. | Epithelial cell marker |
| CD45 | Essential regulator of T- and B-cell antigen receptor signaling. | Hematopoietic cell marker |
| CD68 | Transmembrane glycoprotein and scavenger receptor, expressed by monocytes and macrophages. | Monocyte and tissue macrophage marker |
| PAX8 | Transcription factor. Involved in the development of organs derived from the Mullerian duct. | Secretory epithelial cell marker |
| FOXJ1 | Transcription factor that is involved in the transcription of genes that control cilia production. | Ciliated cell marker |

Figure S3

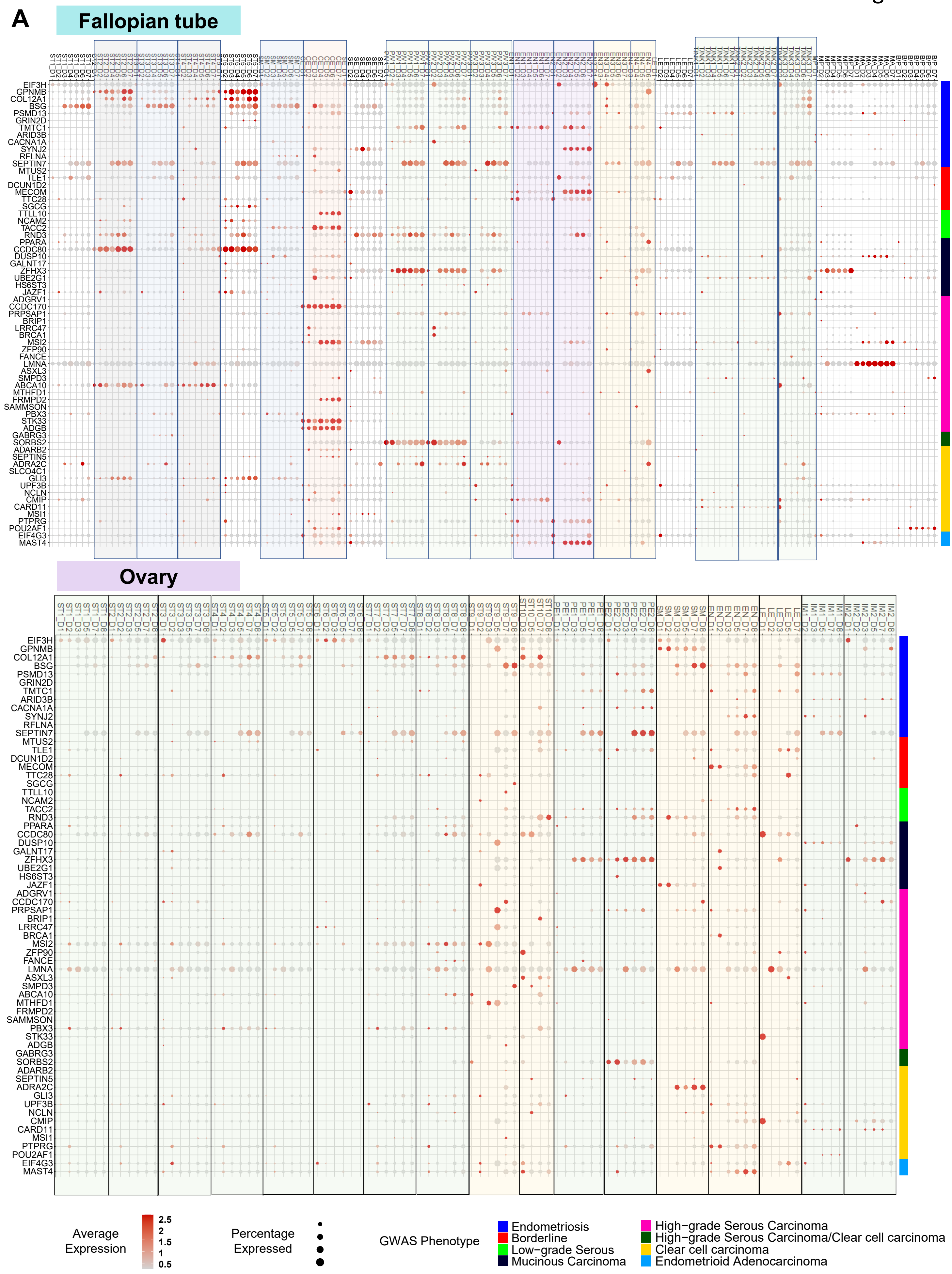

A
Fallopian tube
Ovary
EIF3H
GPNMB
COL12A1
BSG
PSMD13
GRIN2D
TMTC1
ARID3B
CACNA1A
SYNJ2
RFLNA
SEPTIN7
MTUS2
TLE1
DCUN1D2
MECOM
TTC28
SGCG
TTLL10
NCAM2
TACC2
RND3
PPARA
CCDC80
DUSP10
GALNT17
ZFHX3
UBE2G1
HS6ST3
JAZF1
ADGRV1
CCDC170
PRPSAP1
BRIP1
LRRC47
BRCA1
MSI2
ZFP90
FANCE
LMNA
ASXL3
SMPD3
ABCA10
MTHFD1
FRMPD2
SAMMSON
PBX3
STK33
ADGB
GABRG3
SORBS2
ADARB2
SEPTIN5
ADRA2C
SLCO4C1
GLI3
UPF3B
NCLN
CMIP
CARD11
MSI1
PTPRG
POU2AF1
EIF4G3
MAST4
Average Expression
2.5
2
1.5
1
0.5
Percentage Expressed
GWAS Phenotype
Endometriosis
Borderline
Low−grade Serous
Mucinous Carcinoma
High−grade Serous Carcinoma
High−grade Serous Carcinoma/Clear cell carcinoma
Clear cell carcinoma
Endometrioid Adenocarcinoma

Figure S3

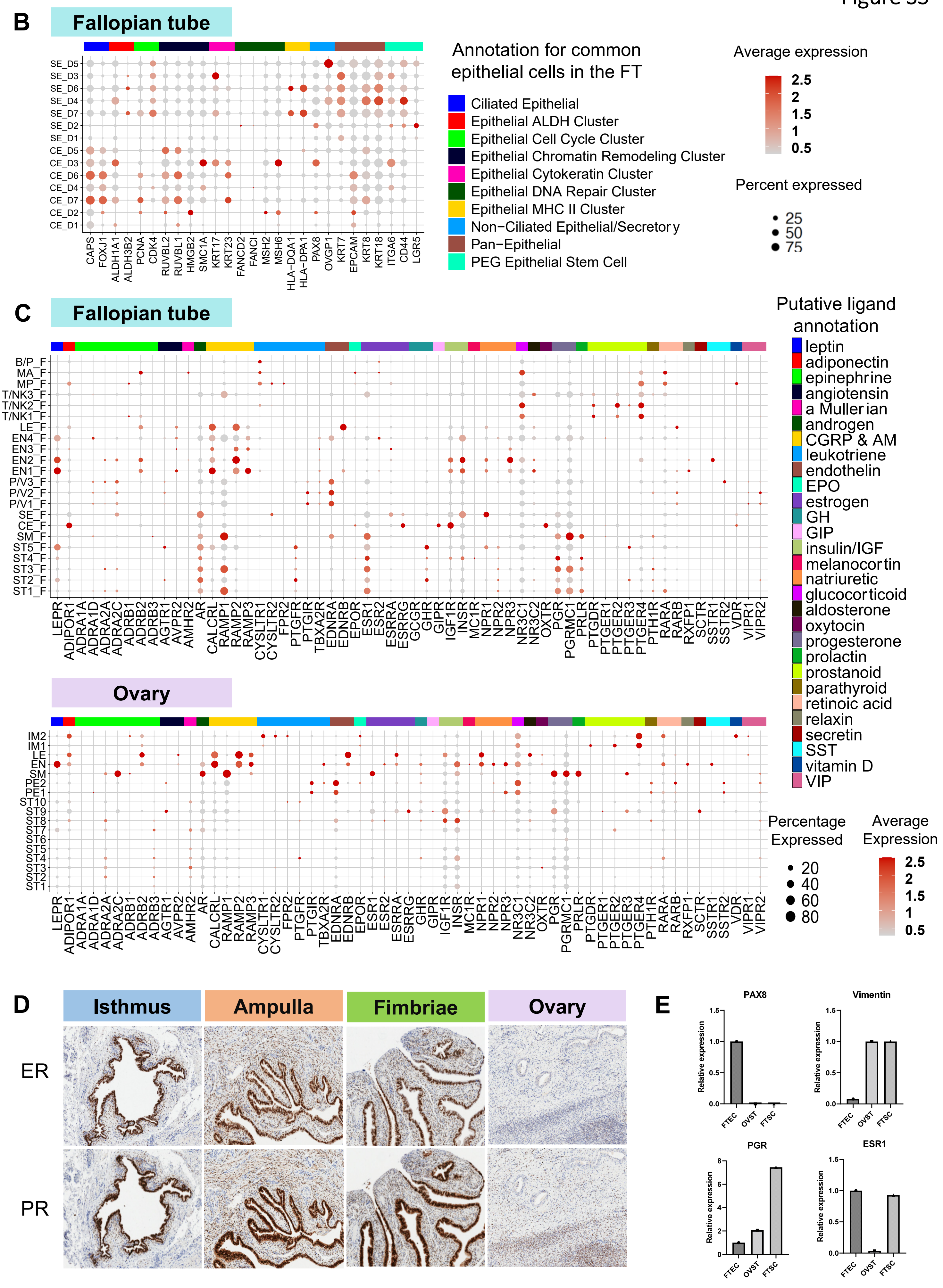

B
Fallopian tube
SE_D5
SE_D3
SE_D6
SE_D4
SE_D7
SE_D2
SE_D1
CE_D5
CE_D3
CE_D6
CE_D4
CE_D7
CE_D2
CE_D1
CAPS FOXJ1 ALDH1A1 ALDH3B2 PCNA CDK4 RUVBL2 RUVBL1 HMGB2 SMC1A KRT17 KRT23 FANCD2 FANCI MSH2 MSH6 HLA-DQA1 HLA-DPA1 PAX8 OVGP1 KRT7 EPCAM KRT8 KRT18 ITGA6 CD44 LGR5
Annotation for common epithelial cells in the FT
Ciliated Epithelial
Epithelial ALDH Cluster
Epithelial Cell Cycle Cluster
Epithelial Chromatin Remodeling Cluster
Epithelial Cytokeratin Cluster
Epithelial DNA Repair Cluster
Epithelial MHC II Cluster
Non−Ciliated Epithelial/Secretory
Pan−Epithelial
PEG Epithelial Stem Cell
Average expression
2.5
2
1.5
1
0.5
Percent expressed
25
50
75
C
Fallopian tube
B/P_F
MA_F
MP_F
T/NK3_F
T/NK2_F
T/NK1_F
LE_F
EN4_F
EN3_F
EN2_F
EN1_F
P/V3_F
P/V2_F
P/V1_F
SE_F
CE_F
SM_F
ST5_F
ST4_F
ST3_F
ST2_F
ST1_F
LEPR ADIPOR1 ADRA1A ADRA1D ADRA2A ADRA2C ADRB1 ADRB2 ADRB3 AGTR1 AVPR2 AMHR2 AR CALCRL RAMP1 RAMP2 RAMP3 CYSLTR1 CYSLTR2 FPR2 PTGFR PTGIR TBXA2R EDNRA EDNRB EPOR ESR1 ESR2 ESRRA ESRRG GCGR GHR GIPR IGF1R INSR MC1R NPR1 NPR2 NPR3 NR3C1 NR3C2 OXTR PGR PGRMC1 PRLR PTGDR PTGER1 PTGER2 PTGER3 PTGER4 PTH1R RARA RARB RXFP1 SCTR SSTR1 SSTR2 VDR VIPR1 VIPR2
Putative ligand annotation
leptin
adiponectin
epinephrine
angiotensin
a Mullerian
androgen
CGRP & AM
leukotriene
endothelin
EPO
estrogen
GH
GIP
insulin/IGF
melanocortin
natriuretic
glucocorticoid
aldosterone
oxytocin
progesterone
prolactin
prostanoid
parathyroid
retinoic acid
relaxin
secretin
SST
vitamin D
VIP
Ovary
IM2
IM1
LE
EN
SM
PE2
PE1
ST10
ST9
ST8
ST7
ST6
ST5
ST4
ST3
ST2
ST1
LEPR ADIPOR1 ADRA1A ADRA1D ADRA2A ADRA2C ADRB1 ADRB2 ADRB3 AGTR1 AVPR2 AMHR2 AR CALCRL RAMP1 RAMP2 RAMP3 CYSLTR1 CYSLTR2 FPR2 PTGFR PTGIR TBXA2R EDNRA EDNRB EPOR ESR1 ESR2 ESRRA ESRRG GCGR GHR GIPR IGF1R INSR MC1R NPR1 NPR2 NPR3 NR3C1 NR3C2 OXTR PGR PGRMC1 PRLR PTGDR PTGER1 PTGER2 PTGER3 PTGER4 PTH1R RARA RARB RXFP1 SCTR SSTR1 SSTR2 VDR VIPR1 VIPR2
Percentage Expressed
20
40
60
80
Average Expression
2.5
2
1.5
1
0.5
D
Isthmus
Ampulla
Fimbriae
Ovary
ER
PR
E
PAX8
Vimentin
PGR
ESR1
Relative expression
FTEC
OVST
FTSC

Figure S4

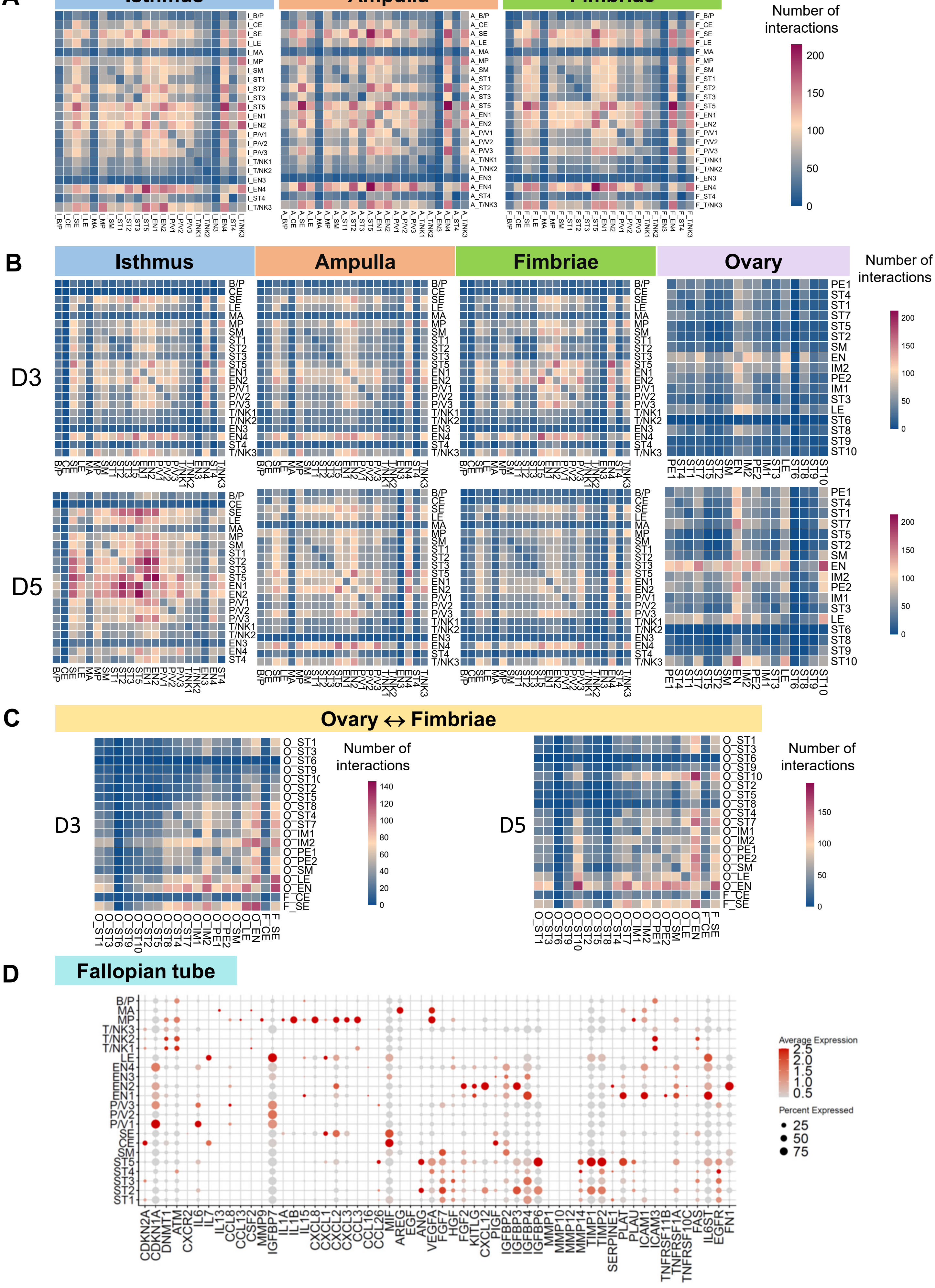
A
Isthmus
Ampulla
Fimbriae
Number of interactions
200
150
100
50
0
B
Isthmus
Ampulla
Fimbriae
Ovary
Number of interactions
D3
D5
C
Ovary ↔ Fimbriae
D3
D5
Number of interactions
D
Fallopian tube
Average Expression
2.5
2.0
1.5
1.0
0.5
Percent Expressed
25
50
75
CDKN2A
CDKN1A
DNMT1
ATM
CXCR2
IL6
IL7
IL13
CCL8
CCL13
CSF2
MMP9
IGFBP7
IL1A
IL1B
IL15
CXCL8
CXCL1
CXCL2
CXCL3
CCL3
CCL16
CCL26
MIF
AREG
EGF
ANG
VEGFA
FGF7
HGF
FGF2
KITLG
CXCL12
PIGF
IGFBP2
IGFBP3
IGFBP4
IGFBP6
MMP1
MMP10
MMP12
MMP14
TIMP1
TIMP2
SERPINE1
PLAT
PLAU
ICAM1
ICAM3
TNFRSF11B
TNFRSF1A
TNFRSF10C
FAS
IL6ST
EGFR
FN1

Figure S5

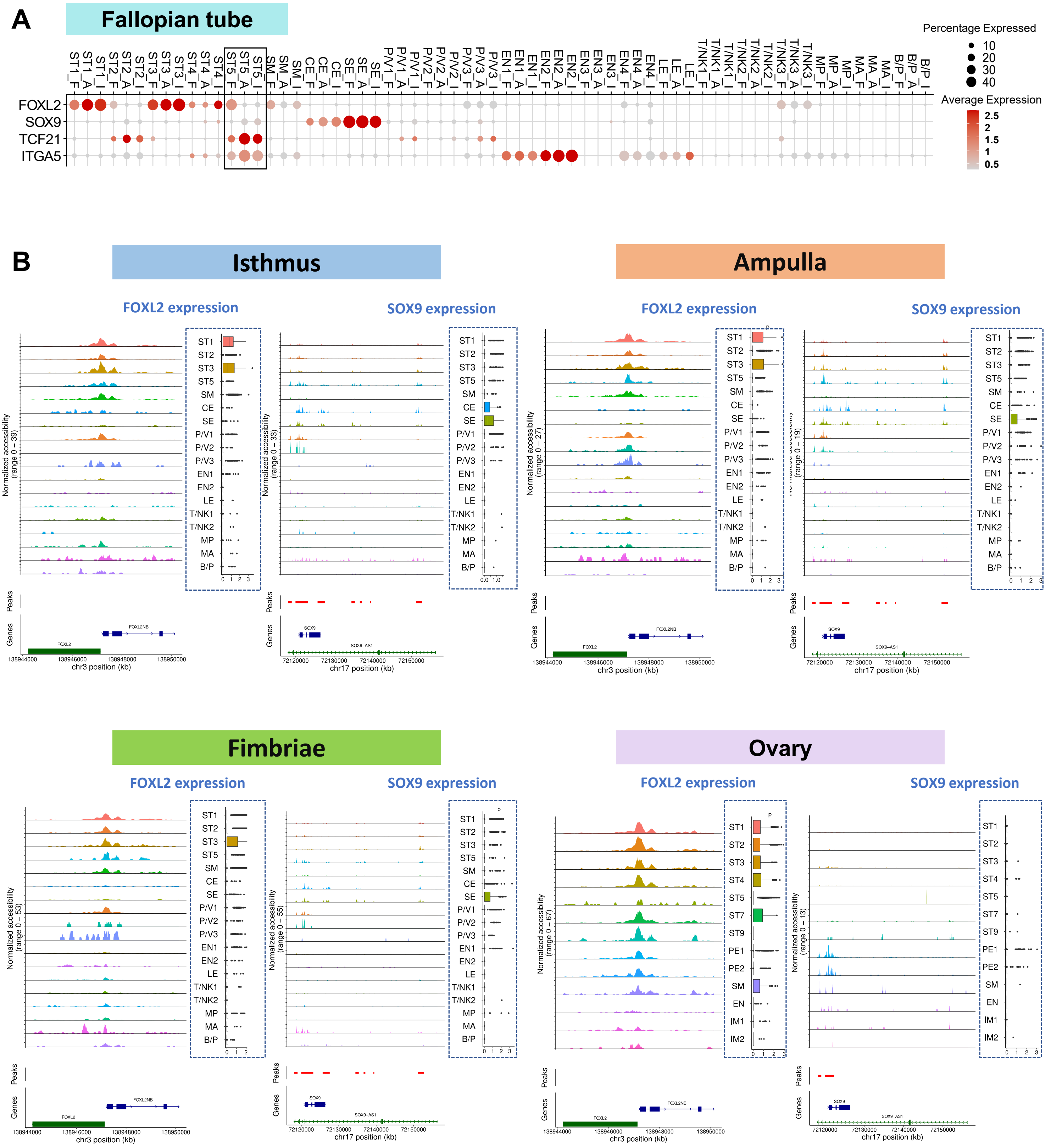
A
Fallopian tube
Percentage Expressed
Average Expression
FOXL2
SOX9
TCF21
ITGA5
B
Isthmus
Ampulla
Fimbriae
Ovary
FOXL2 expression
SOX9 expression
Normalized accessibility
Peaks
Genes
chr3 position (kb)
chr17 position (kb)

A

Figure S6

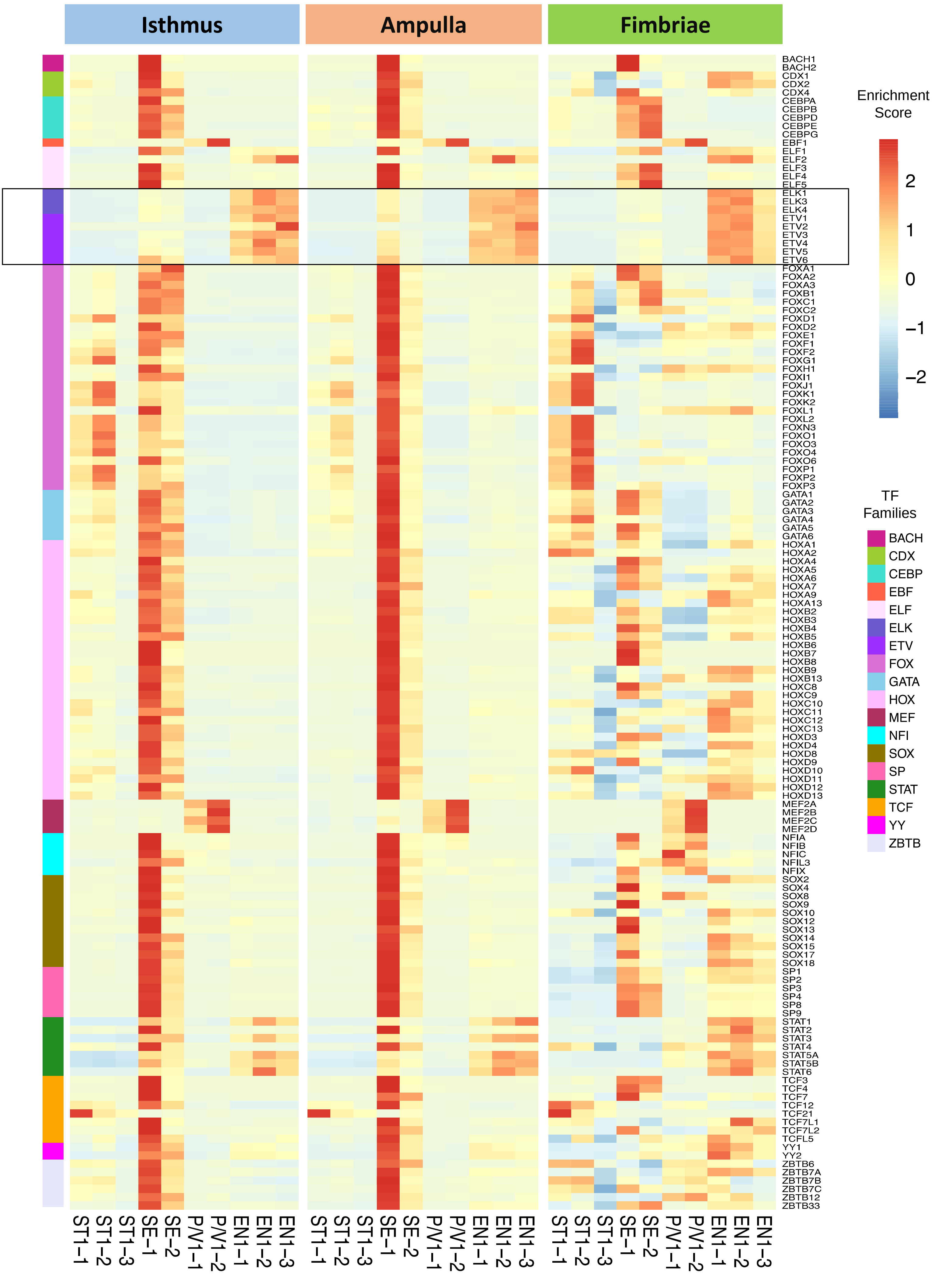

Isthmus
Ampulla
Fimbriae
Enrichment Score
2
1
0
−1
−2
TF Families
BACH
CDX
CEBP
EBF
ELF
ELK
ETV
FOX
GATA
HOX
MEF
NFI
SOX
SP
STAT
TCF
YY
ZBTB
ST1−1
ST1−2
ST1−3
SE−1
SE−2
P/V1−1
P/V1−2
EN1−1
EN1−2
EN1−3

**B**

Figure S6

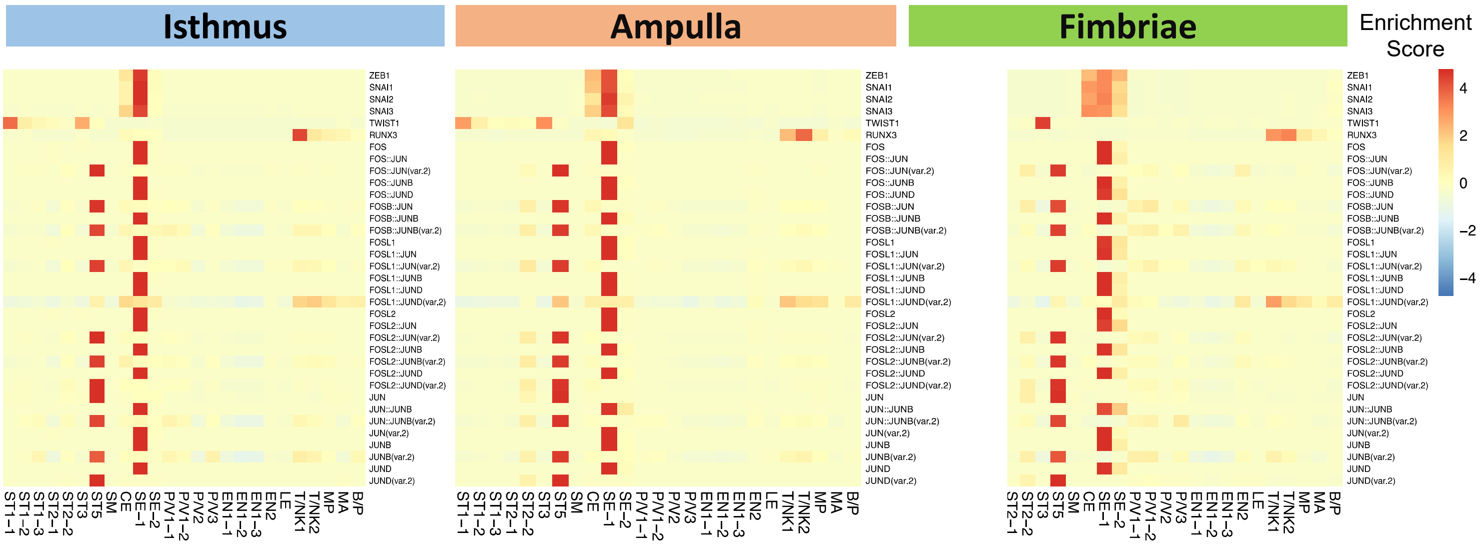

Isthmus
Ampulla
Fimbriae
Enrichment Score
4
2
0
−2
−4
ZEB1
SNAI1
SNAI2
SNAI3
TWIST1
RUNX3
FOS
FOS::JUN
FOS::JUN(var.2)
FOS::JUNB
FOS::JUND
FOSB::JUN
FOSB::JUNB
FOSB::JUNB(var.2)
FOSL1
FOSL1::JUN
FOSL1::JUN(var.2)
FOSL1::JUNB
FOSL1::JUND
FOSL1::JUND(var.2)
FOSL2
FOSL2::JUN
FOSL2::JUN(var.2)
FOSL2::JUNB
FOSL2::JUNB(var.2)
FOSL2::JUND
FOSL2::JUND(var.2)
JUN
JUN::JUNB
JUN::JUNB(var.2)
JUN(var.2)
JUNB
JUNB(var.2)
JUND
JUND(var.2)
ST1−1
ST1−2
ST1−3
ST2−1
ST2−2
ST3
ST5
SM
CE
SE−1
SE−2
P/V1−1
P/V1−2
P/V2
P/V3
EN1−1
EN1−2
EN1−3
EN2
LE
T/NK1
T/NK2
MP
MA
B/P

### Supplementary Figure Legends

#### Figure S1 (related to Figure 1). Mapping major cell types of the normal human postmenopausal fallopian tube and ovary.

A) Dot plot showing normalized gene expression levels of known canonical marker genes for each cell type identified in the fallopian tube in Fig. 1B.

B) Scatter plot comparing gene expression levels of different stromal cell populations: ST2/5 *versus* ST1/3/4.

C) Canonical cell types in the normal human postmenopausal fallopian tube. UMAP plot showing the 11 major cell clusters identified in the fallopian tube using scRNA-seq. Data includes all three anatomic regions (Fimbriae, ampulla, and isthmus) for a total of seven donors. Relative abundance of the 11 major cell clusters found in the postmenopausal fallopian tube using scRNA-seq. The graph shows the individual percentage of each cell type by the individual donor.

D) Dot plot showing normalized gene expression levels of known canonical marker genes for each cell type identified in the postmenopausal ovary in Fig. 1D.

E) Canonical cell types in the normal human postmenopausal ovary. UMAP plot showing the six major cell clusters identified in the ovary using scRNA-seq. Relative abundance of the 11 major cell clusters found in the postmenopausal ovary using scRNA-seq. Data includes a total of six patients. The graph shows the individual percentage of each cell type by the individual donor.

F) Hematoxylin and eosin (H&E) staining and RNA-fluorescent in situ hybridization (FISH) of decorin (DCN) in the postmenopausal ovary from donor D3. Every dot corresponds to one RNA transcript. Nuclei are stained with DAPI in cyan and DCN transcripts in yellow. Dashed white lines separate the stroma from blood vessels. Scale bar is 100 µm.

G) Dot plot showing average gene expression of markers associated with oocyte and follicle development in the fallopian tube and ovary.

H) Fallopian tube, scRNA-seq: Comparison of our scRNA-seq data from the fallopian tube of 7 postmenopausal women (noted as ‘HCA’), with normal epithelial cells from Hu, et al., Cancer cell 2020, and one postmenopausal patient from Dinh, et al., Cell Reports, 2021. Cells from the three datasets were integrated, clustered, and annotated based on our annotations. J) UMAPs comparing two published datasets with our data in the same UMAP space. Three new clusters named ‘NEW1/2/3’ appear in ‘Cancer Cell, 2020’ data only; cell types identified in one postmenopausal patient from ‘Cell Reports, 2021’ showed good overlap with cell types identified by us.

Abbreviations: ST= stromal cells, T/NK= T cells and NK cells, SE= secretory epithelial cells, LE= lymphatic endothelial cells, SM= smooth muscle cells, MP= macrophages, P/V= pericytes and vascular smooth muscle cells, CE= ciliated epithelial cells, EN= endothelial cells, B/P= B cells and plasma B cells, MA= mast cells, NEW1-3= Unknown clusters detected from Cancer cell 2020 dataset only.

### Figure S2 (related to Figure 2). Expression profiles of the isthmus, ampulla, and fimbrial regions of the postmenopausal fallopian tube.

A) Immunohistochemical staining of vimentin, CD45, CD68 and EPCAM in the postmenopausal isthmus, ampulla, fimbriae, and ovary. PAX8 and FOXJ1 expressions are shown in the isthmus, ampulla, and fimbriae of the fallopian tube (scale bar 100 μm).

### Figure S3 (related to Figure 3). Expression of GWAS and other marker genes in different regions of the postmenopausal fallopian tube and the ovary obtained from scRNA-seq.

A) Cell type specific GWAS gene expression in the fallopian tube and ovary in different patients. Fallopian tube (top): Cell subtypes with matching colors (grey, blue, green, and purple) have that same set of GWAS genes expressed *among* and *across* subtypes, except for yellow where some patients express different sets of GWAS genes. Ovary (bottom): Cell subtypes highlighted in green show expression for the same set of GWAS genes among patients, and in yellow where some patients express different sets of GWAS genes.

B) Normalized expression of genes associated with ciliated (CE) and secretory (SE) epithelial cells of the fallopian tube in each donor (scRNA-seq). The genes are grouped by epithelial cell subtypes or functions. Note the expression of markers for intercalated or PEG cells in SE cells.

C) Dot plots of hormone receptor genes and their putative hormone ligands (color bars on top) in the postmenopausal fallopian tube fimbriae and ovary. Colors indicate the specific hormones.

D) Immunohistochemical staining of estrogen (ER) and progesterone hormone receptors (PR) in the postmenopausal isthmus, ampulla, fimbriae, and ovary. All images are shown at 20x magnification.

E) QRT-PCR of fallopian tube epithelial cells (FTEC), fallopian tube stromal cells (FTSC) and ovarian stromal cells (OVST). Bar graphs show relative expression of PAX8, Vimentin, PGR and ESR1 for each cell type.

### Figure S4 (related to Figure 4). Ligand-receptor interactions in the fallopian tube and ovary in select patients.

A) Fallopian tube. Heatmaps showing the frequency of interactions among the different cell types in the isthmus, ampulla, and fimbria were detected using CellPhoneDB for all donors. ST5 from the FT interacts strongly with CE, SE, and EN in the FT.

B) Heatmaps showing the frequency of interactions among different cell types in the isthmus, ampulla, and fimbriae in the fallopian tube and ovary in two donors imputed by CellPhoneDB from scRNA-seq.

C) Ovary ⇔ Fimbriae interactions. Heatmaps showing the frequency of interactions detected by CellPhoneDB among different cell types in the ovary (O) with ciliated (CE) and secretory epithelial cells (CE) in the fallopian tube in two donors using scRNA-seq.

D) Aging and senescence-related gene expression in the fallopian tube.

### Figure S5 (related to Figure 5). Analysis of select transcription factors in the postmenopausal fallopian tube and ovary using scRNA-seq and scATAC-seq.

A) Dot plot of select EMT markers in the fallopian tube by the anatomical site (Isthmus: I, ampulla: A, and fimbria: F) based on scRNA-seq.

B) Track plots from scATAC-seq showing the accessibility profiles at FOXL2 and SOX9 loci across all cell sub-types (left) in the isthmus, ampulla, fimbriae, and ovary. Matching scRNA-seq expressions of FOXL2 and SOX9 are shown in the boxplots (right).

### Figure S6 (related to Figure 6). Transcription factor (TF) analysis of scATAC-seq data in the different anatomical regions of the fallopian tube.

A) Heat maps depicting select transcription factor accessibility in the isthmus, ampulla and fimbria by cell type using scATAC-seq alone. Cell types include secretory epithelial SE-1, SE-2, stromal ST1-1, ST1-2, ST1-3, endothelial EN1-1, EN1-2, and pericytes/vascular endothelial P/V1-1, P/V1-2 cells. TF families ELK and ETV are highlighted.

B) TF enrichment of JUN/FOS family and other EMT TFs including ZEB1, SNA12, SNA13, TWIST1 and RUNX3 in the in the isthmus, ampulla and fimbria (using the JASPAR transcription factor database). For a few TF, the splicing variant (var2) is expressed in one cell type but not in the other.
